# In vivo vaccine generation with a topical small molecule cocktail to eradicates AML and PDAC

**DOI:** 10.1101/2025.10.19.683274

**Authors:** Shuwen Zheng, Mengyuan Li, Zhaoxing Wu, Xuzhao Zhang, Xiaoxian Gan, Rongzhen Xu

## Abstract

Using vaccine to cure cancer has been expected for over 130 years, but remains a big challenge. Here we present a novel in vivo personal and off-the-shelf vaccine generation strategy with unique mechanism, which is very simple but high efficacy, using a small molecule cocktail as antigen-agnostic vaccine. In mouse AML and PDAC models, this in vivo vaccine rapidly eliminated tumors via triggering inflamed immunogenic cell death to release antigens, activating NK and DC cells, inducing innate and adaptive immunity in blood, lymph node, spleen, bone marrow and TME, and elicited a robust long-term immune memory. Importantly, this small molecule cocktail-based platform holds the potential to make in vivo vaccine more personal and accessible. It might represent a promising avenue for pan-cancer immunotherapy.

## Introduction

Harnessing the immune system with vaccines to recognize and combat cancer was a concept initially demonstrated by William Coley in the early 1900s. Since then, extensive efforts have attempted to use cancer vaccines to induce antitumor immune responses but with limited clinical success^1^. Etiologies of immunotherapy resistance are multifaceted but largely associated with the deficiency of the cancer-immunity cycle ^2, 3, 4^. As most tumors frequently escape a single immune cell-mediated immunity ^5^, thus, the entire cancer-immunity cycle is crucial for eradicating heterogeneous tumors. The host retains innate and adaptive immune cells for recognizing and killing tumor cells, however, their activity is modulated through stimulatory and inhibitory components^3^. Once tumor is well established, the balance between these inputs is tipped toward immunosuppression ^6, 7^, resulting in immune evasion. Given that most tumors harbor sufficient personal neoantigens for vaccines ^8, 9, 10^, but tumors have an enormous molecular diversity, and selecting the neoantigens for immune recognition is a hugely complex task. Hence, direct in vivo cancer vaccination may be the best solution to release patient’s unique neoantigens and stimulate a robust immune response to clear various tumors if the deficient cancer-immunity cycle can be reversed. We speculate that the successful in vivo vaccine requires three key components: (1) potent triggers of immunogenic cancer cell death (ICD) to release sufficient immunogenic neoantigens, (2) strong activators of DC/APC cells for effective antigen presentation to T cells, and (3) robust enhancers of diverse innate and adaptive immune cells to clear various tumors with immune memory.

By screening small molecule immunomodulators, we have identified several small chemical molecules as promising candidates to activate cancer-immunity cycle. Small molecule resiquimod (RMQ) can trigger an extensive activation of the innate immune cells, including NK cells and DCs^11, 12, 13, 14^, and topical resiquimod can induce tumor regression and enhance T-cell effector functions in human cutaneous T-cell lymphoma (CTCL)^15^. PT100 is a small molecule inhibitor of dipeptidyl peptidase (DPP) and improves T cell infiltration and reverse the immunosuppressive tumor microenvironment, accelerates T cell priming ^16^. Importantly, PT100 alters the CXCR3 axis and enhances NK and CD8+ T cell infiltration in murine models of PDAC ^17^. In addition, PT100 also remodels the tumor microenvironment in Lewis lung carcinoma model ^18^. Dithranol (DTH), an anti-psoriatic small molecule agent, is a strong inflammatory inducer that can boost TLR7 agonist-induced durable CD8+ and CD4+ T cell responses and skin-resident memory responses ^19, 20^. Although monotherapy of these small molecules was of limited therapeutic benefit, we hypothesized that the cocktail of these small molecules with different targets may have synergistic effect and fulfill requirement of key components for in vivo vaccine generation. Moreover, human skin is a potent immune-competent organ ^21^. The non-invasive transdermal administration of small molecules not only can target skin-resident professional antigen-presenting cells (APC) priming high quality T cell responses in draining lymph nodes as well as systemic immune system, but also is well safety and simplified clinical medication.

In this study, we aimed to develop a novel in vivo vaccine generation strategy with high efficacy, safety, and simplicity for pan-cancer immunotherapy using a topical small molecule cocktail formulation as antigen-agnostic cancer vaccine. Using acute myeloid leukemia (AML) and pancreatic ductal adenocarcinoma (PDAC) with resistance to current immunotherapies as models ^22, 23, 24, 25, 26, 27, 28^, we conducted the comprehensive proof-of-concept in vitro and in vivo to identify its target immune cells and mechanisms of action. Unlike current cancer vaccines that predominantly boost CD8^+^ T cells, this novel in vivo vaccine can potently and extensively activate both innate and adaptive immunity with long-term memory and generates a super pan-immune cell network to clear multiple tumors.

## Results

### Topical small molecule cocktail-based in vivo vaccine eradicates acute myeloid leukemia with long-term immune memory

In order for this technology to be used broadly in cancer immunotherapy with an acceptable safety, we sought to develop an in vivo vaccine via non-invasive topical administration with a small molecule cocktail as antigen-agnostic cancer vaccine. The small molecule cocktail-based vaccine formulation consists of RMQ, PT100 and DTH. We rationally designed RMQ and PT100 as super-activators of NK and DC cells to trigger immunogenic cell death (ICD) and release tumor antigens, and promote process and present tumor antigens, which systemically activate innate and adaptive immunity to clear various tumors. Importantly, we designed DTH as an inflammatory adjuvant to improve in vivo vaccine-induced immune responses and long-term memory. In addition, we developed an all-in-one formulation with Vaseline cream, which can be easily used for patients.

We next investigated the efficacy and safety of this topical small molecule cocktail-based in vivo vaccine or its key components RMQ or PT100 using AML, which is highly resistance to current therapies ^22, 23, 24, 25^, as a model. We first established a systemic murine AML model using C57BL/6 mice with murine AML C1498 cells, which is a highly aggressive disseminated AML model originally isolated from a leukemic 10-month-old C57BL/6 (H-2b) female mouse.

C57BL/6 mice were inoculated with 5 × 10^6^ luciferase-expressing C1498 cells through tail vein. After the tumor signal was detected, mice were randomly divided into four groups to receive vehicle or RMQ or PT100 or the cocktail via topical application once a week for 3 weeks. As shown in Fig. 1, single component alone treatment resulted in transient suppression of established leukemia in mice and the majority of these mice subsequently succumbed to disease (Fig. 1A, B). By contrast, the cocktail treatment increased the proportion of responding mice to 100%, and induced a durable complete remission of tumor growth after receiving a 3-week period of transdermal application (Fig. 1C, D, E). The TGI values of RMQ, PT100 and cocktail were 69.81%, 47.84% and 97.89%, respectively (Fig. 1F). Consistently, the cocktail treatment for 3 weeks significantly improved the median and overall survival of AML mice. However, untreated mice quickly succumbed to death with a median survival time (MST) of only 20 days due to the aggressive disease progression and high tumor burden (Fig. 1G).

**Fig. 1.**
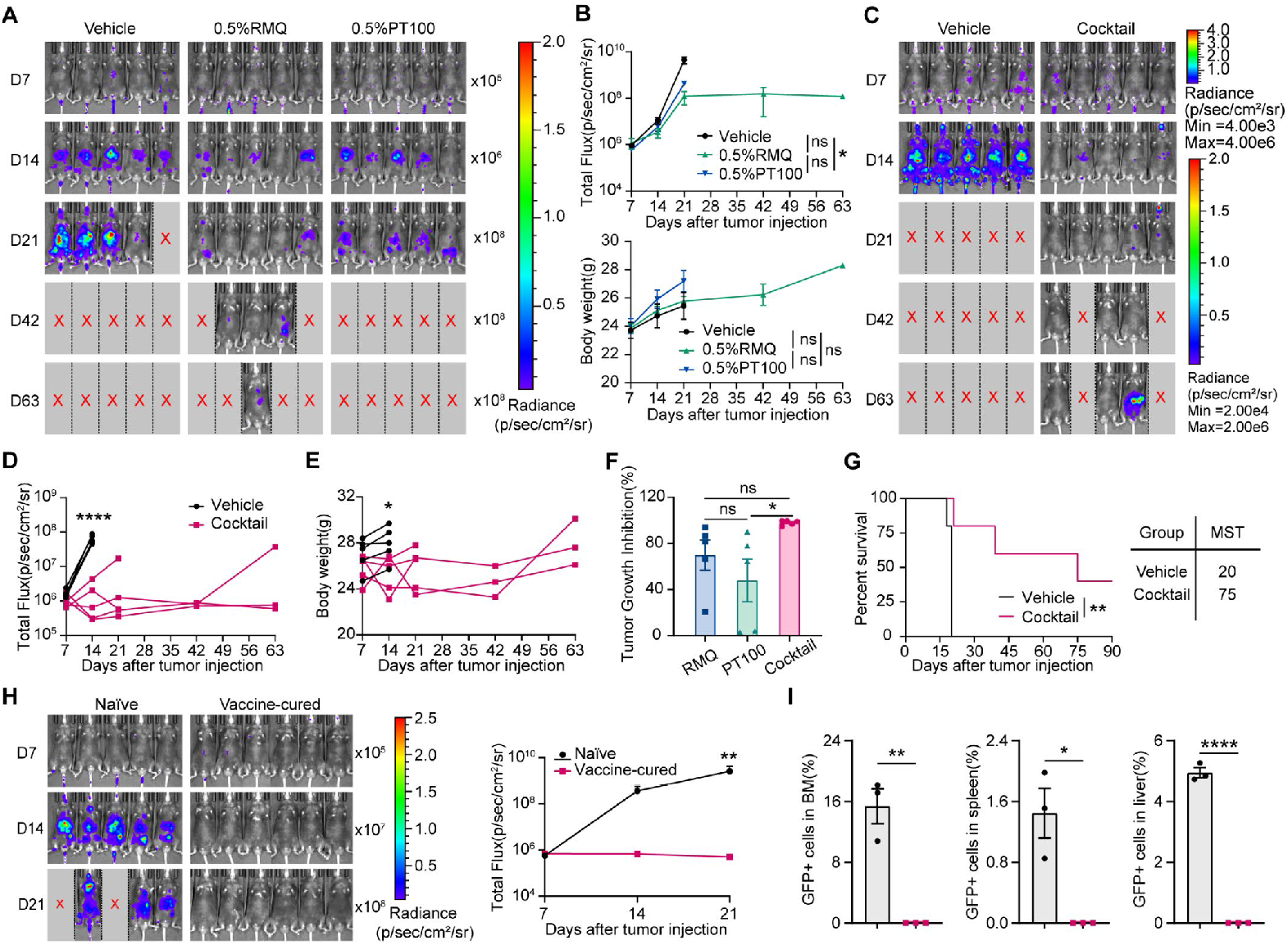
Topical small molecule cocktail-based in vivo vaccine eradicates acute myeloid leukemia with long-term immune memory in mouse Model. **A**. Bioluminescence imaging of C1498 xenograft C57 mice treated with vehicle, 0.5%RMQ or 0.5%PT100 (n=5). **B**. Quantification of tumor burden and body weight of experimental mice at the indicated days. **C**. Bioluminescence imaging of C1498 xenograft C57 mice treated with vehicle or cocktail (n=5). **D. E**. Quantification of tumor burden (**D**) and body weight (**E**) of experimental mice at the indicated days. **F**. Tumor Growth inhibition (TGI) of mice treated with 0.5%RMQ, 0.5%PT100 or cocktail (n=5). **G**. Kaplan-Meier plot showing mouse survival. **H**. Bioluminescence imaging and quantification of tumor burden of C1498 leukemia cells rechallenged C57 naïve (n=5) and vaccine-cured mice (n=6). **I**. Representative flow cytometry graphs and calculated percentage of C1498-GFP cells in mice bone marrow (BM), Spleen and Liver. Gating strategy is depicted in **Fig. S2E**. All error bars represent means ± s.e.m. Statistical significance was determined by one-way ANOVA and two-tailed t-test. * p<0.05, ** p<0.01, **** p<0.0001, ns means no significance.

Of note, 100% of mice treated with the cocktail survived for over 28 days and 60% mice (3/5) were tumor free in AML murine model. Moreover, 40% mice (3/5) remain tumor free at day 90 after AML C1498 transplantation (Fig. 1G). These results indicate that anti-tumor immunity induced by single component rapidly fade away, resulting in poor memory formation, whereas the cocktail-based in vivo vaccine confers deep and durable complete remission of AML in mouse model.

To evaluate the safety of the cocktail-based in vivo vaccine, we measured the body weight of mice, and found that topical cocktail treatment led to a transient weight loss of 2.0% compared to control animals at 7 days after receiving the first dose of topical cocktail, however, the weight loss was fully reversible after completing the treatment cycle (Fig. 1E). Moreover, no adverse effects were observed at treated skin sites with cocktail or single component RQM or PT100.

These results suggest that the topical cocktail-based in vivo vaccine is safe for use.

Given the potent efficacy observed in mouse AML model, we next asked whether the in vivo vaccine can induce a long-term anti-tumor immune memory. We rechallenged in vivo vaccine-cured mice with 5×10^6^ C1498-luciferase-GFP leukemia cells through tail vein after initial leukemia cell transplantation for 90 days. Naïve C57BL/6 mice inoculated with C1498-luciferase-GFP leukemia cells served as control. We found that that all in vivo vaccine-cured mice resisted subsequent leukemia cell rechallenge, whereas leukemia cells grew robustly in naïve mice (Fig. 1H). All naïve mice were sacrificed at 21 days post-re-challenge. A substantial number of leukemia cells were found in BM (15.4%), spleen (1.4%), liver (5.0%) of naïve mice (Fig. 1I), that implying long-lasting protective immunity.

### Small molecule cocktail-based in vivo vaccine eliminates both local and distant PDAC tumors with immune memory

The superior therapeutic effect of the cocktail-based in vivo vaccine against AML encouraged us to further evaluate the therapeutic effect on the established solid tumor PDAC, which has the characteristics of high-density desmoplastic stroma, a distinctive immunosuppressive microenvironment and is profoundly resistant to all forms of chemotherapy and immunotherapy ^26, 27, 28^.

To investigate whether the topical in vivo vaccine inhibits local and distant solid tumor growth, we used the KPC syngeneic mouse model for pancreatic cancer. We first established two-separate-tumor bearing mouse PDAC models. Luciferase-expressing KPC cells were injected subcutaneously into syngeneic C57BL/6 mice at left (local) and right (distant) sites. When the largest diameter of tumor reached between 0.5 and 0.8 cm, the cocktail or vehicle was topically applied on the left tumor site (local) of mice for once a week for 3 weeks. Tumor growth at both left (local) and right (distant) sites and animal body weight were then recorded (Fig. 2). The tumors of vehicle-treated mice grew rapidly at both left and right sites, until it reached the human endpoint within 28 days (Fig. 2A-C). However, the cocktail treatment caused a complete regression of both local and distant tumors among nearly all of the mice (Fig. 2A-D). Cocktail-treated local tumor regressed completely within 7 days of the first dose. Distant tumor (untreated) regression was slightly delayed, but tumors were also completely eliminated after the second dose. All these results indicate that the cocktail-based in vivo vaccine has excellent therapeutic effect in the mouse PDAC model.

**Fig. 2.**
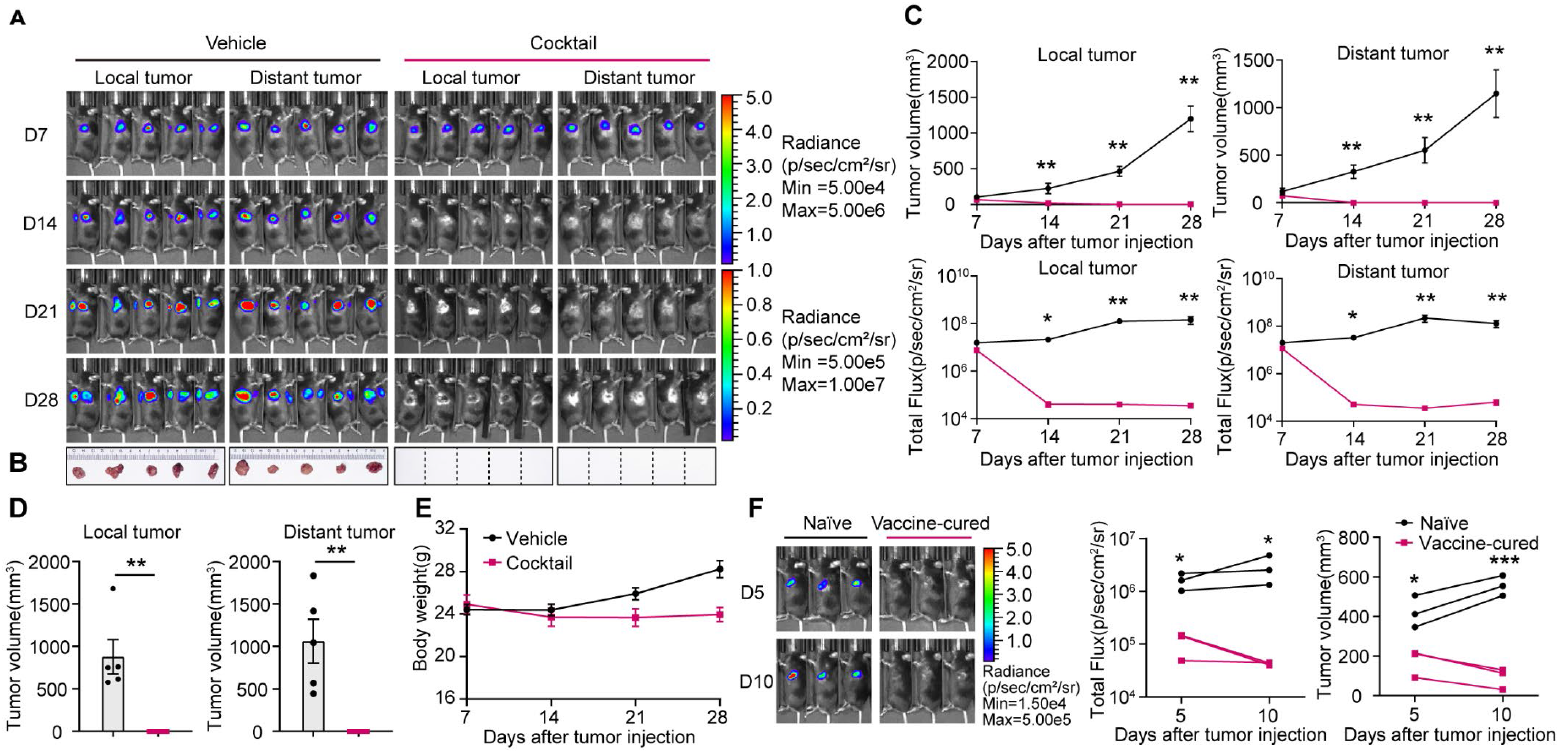
Small molecule cocktail-based in vivo vaccine eliminates both local and distant PDAC tumors with immune memory. **A**. Bioluminescence imaging of KPC xenograft C57 mice treated with vehicle or cocktail (n=5). **B**. Gross appearances of tumors after treatment with vehicle or cocktail for 28days (n=5). **C. D. E**. Local and distant tumor volume and quantification of tumor burden (**C. D**) of KPC xenografts and mouse weight (**E**) after treatment with vehicle or cocktail at indicated time. **F**. Bioluminescence imaging and quantification of tumor loads and tumor volume of KPC rechallenged C57 naïve and vaccine-cured mice (n=3). All error bars represent means ± s.e.m. Statistical significance was determined by the two-tailed t-test. * p<0.05, ** p<0.01, *** p<0.001.

Consistent with mouse leukemia models, the topical cocktail-based in vivo vaccine treatment resulted in a transient weight loss of 4.90% compared to control animals at 7 days after receiving the first dose of transdermal cocktail in mouse PDAC models, but the weight loss was fully reversible after one treatment cycle (Fig. 2E).

To verify whether the in vivo vaccine-cured mice have developed long lasting tumor-specific immunity, they were re-challenged with a same dose of KPC cells. Consistently, all of these animals exhibit tumor-free survival, whereas naïve controls perished. These results further confirm that the cocktail-based in vivo vaccine also elicits long-term immune memory for solid tumor PDAC (Fig. 2F).

### The in vivo vaccine triggers inflamed immunogenic cell death (ICD) with long-term memory

Cancer cells harbor sufficient neoantigens and inflamed immunogenic cell death (ICD) can trigger these neoantigen release and elicit immune response and memory ^8, 9^. To assess whether the cocktail-based in vivo vaccine can induce inflamed immunogenic cell death, we performed histopathological examination and analyzed intratumoral inflammatory immune cell profile in both local (treated) tumor and distant (untreated) tumor tissues from mice at day 7 after one dose of topical cocktail treatment. Tumor tissues were processed for H&E staining, immunostaining of myeloid inflammatory phenotype CD11b^+^/Ly6G^+^/Ly6C^+^ cells and TUNEL staining for apoptosis. We found that one dose of in vivo vaccine treatment caused significant shrinkage of both local and distant tumors in mouse models, showing inflammatory necrosis of tumor tissues with marked increases of CD11b^+^/Ly6G^+^/Ly6C^+^ cells (Fig. 3A-E). Inflammatory CD11b^+^, Ly6G^+^, Ly6C^+^ cell populations increased by 9.02-, 14.50-, and 7.63-folds in TME (Fig. 3C-E), respectively. Conversely, tumor tissues from vehicle-treated mice displayed histological features of tumor with high proliferative activity. Tumor shrinkage and necrosis were positively correlated with inflammatory cells (Fig. 3A-E). These observations indicate that cocktail-based in vivo vaccine induces robust inflamed immunogenic cell death.

**Figure 3.**
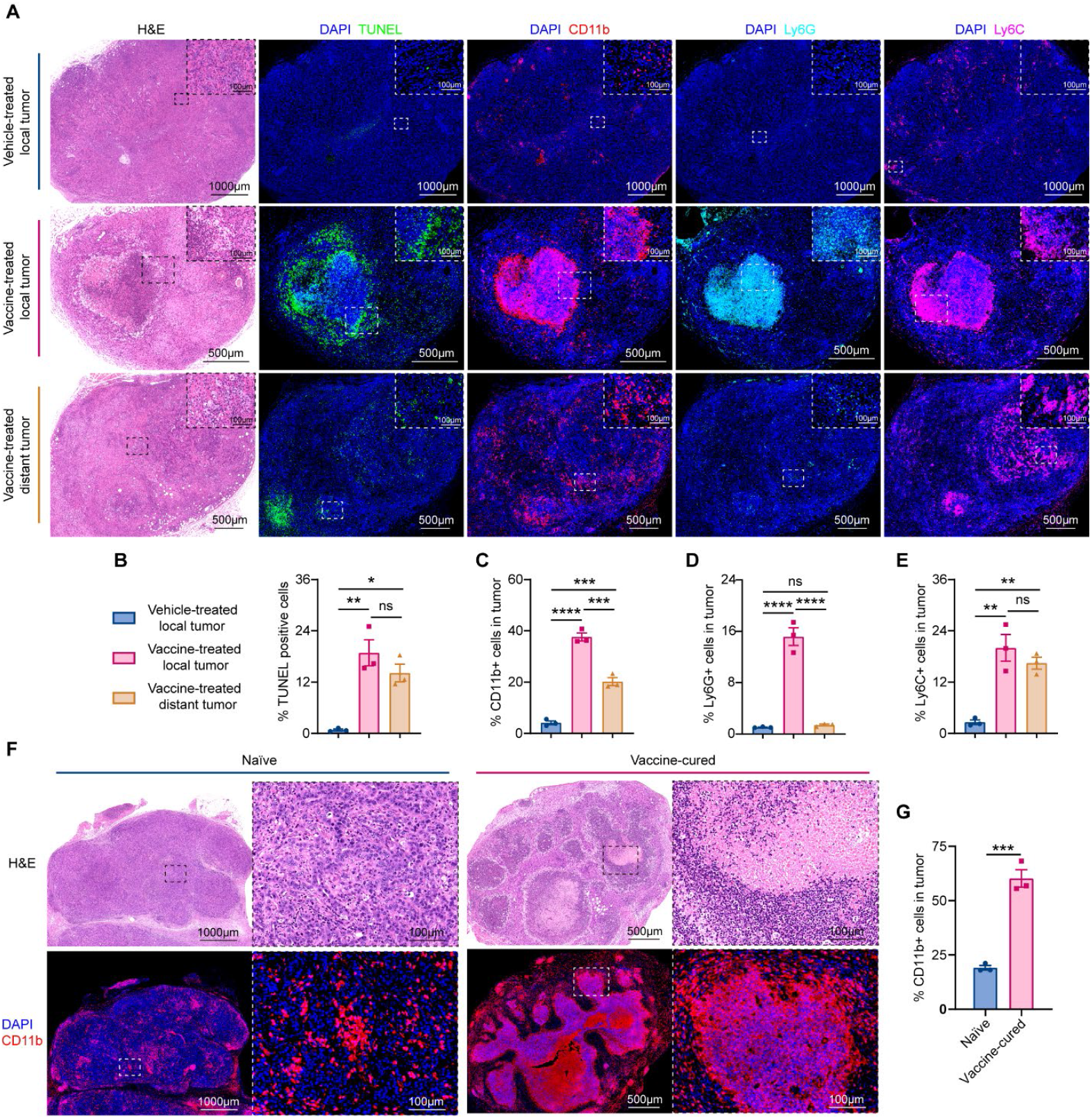
The In vivo vaccine initiates inflamed immunogenic cell death (ICD) in tumor tissues and rechallenged tumor tissues. **A**. Representative and the corresponding local enlarged images of H&E, TUNEL (Green), CD11b (Red), Ly6G (Cyan), Ly6C (Magenta) staining of mice tumor from each treatment group, counterstained with DAPI (Blue). **B-E**. Quantification of TUNEL positive (**B**), CD11b^+^ (**C**), Ly6G^+^ (**D**), Ly6C^+^ (**E**) cells in tumor tissue (n=3). **F**. Representative (left) and the corresponding local enlarged (right) images of H&E and CD11b (Red) staining of mice tumor from naïve and vaccine-cured mice tumor, counterstained with DAPI (Blue). **G**. Quantification of CD11b^+^ cells in tumor tissue (n=3). All error bars represent means ± s.e.m. Statistical significance was determined by one-way ANOVA. * p<0.05, ** p<0.01, *** p<0.001, **** p<0.0001, ns means no significance.

To further confirm above observations, we rechallenged in vivo vaccine-cured mouse PDAC model after initial KPC-luciferase PDAC cells transplantation for 90 days with the same KPC-luciferase PDAC cells. 2×10^6^ Luciferase-expressing KPC cells were injected subcutaneously into vaccine-cured mouse PDAC model at left site. Naïve C57BL/6 mice inoculated with KPC-luciferase PDAC cells served as control. After 7 days of engraftment, implanted tumors were collected for H&E staining and CD11b^+^ myeloid inflammatory cell immunostaining. Consistent with above findings, the implanted tumors showed significant necrosis with concomitant increase of CD11b^+^ cells in vaccine-cured mouse PDAC model, whereas implanted tumors grew robustly in naïve mice (Fig. 3F). Inflammatory CD11b^+^ cells increased by 3.15-folds in TME (Fig. 3G). These findings further support that the in vivo vaccine not only induces potent immunogenic cell death but also elicit long-term immune memory.

### The in vivo vaccine expands NK cells in the blood, spleen, and bone marrow in tumor-bearing mice

We have observed that the in vivo vaccine can clear circulating leukemia cells and solid tumor, and induces a transient mild weight loss following the first dose of topical cocktail in mouse models. These findings suggest that the in vivo vaccine may lead to systemic activation of immunity. Given that NK cells are first-line defenders of the immune response against cancer, however, cancer patients, such as AML and PDAC, frequently exhibits impaired NK cell functions, which can facilitate escape from immune surveillance^29, 30, 31, 32, 33, 34^. To evaluate whether the in vivo vaccine can stimulate NK cells in mouse models, we next measured NK cell frequencies in circulating blood samples of vehicle- and vaccine-treated mouse AML and PDAC models after 7 days of treatment with flow cytometry (Fig. 4). The overall frequency of NK cells significantly increased in the circulating bloods in both cocktail-treated mouse AML and PDAC models (Fig. 4A, B). In mouse AML model, the frequency of NK cells increased by 2.70-fold following vaccine treatment (Fig. 4A) (2.69% NK cell (vehicle) vs. 7.26% NK cell (vaccine), P < 0.0001). Similarly, the frequency of NK cells increased by 3.72-fold following vaccine treatment (Fig. 4B) (1.86% NK cell (vehicle) vs. 6.91% NK cell (vaccine), P < 0.0001) in mouse PDAC model.

**Figure 4.**
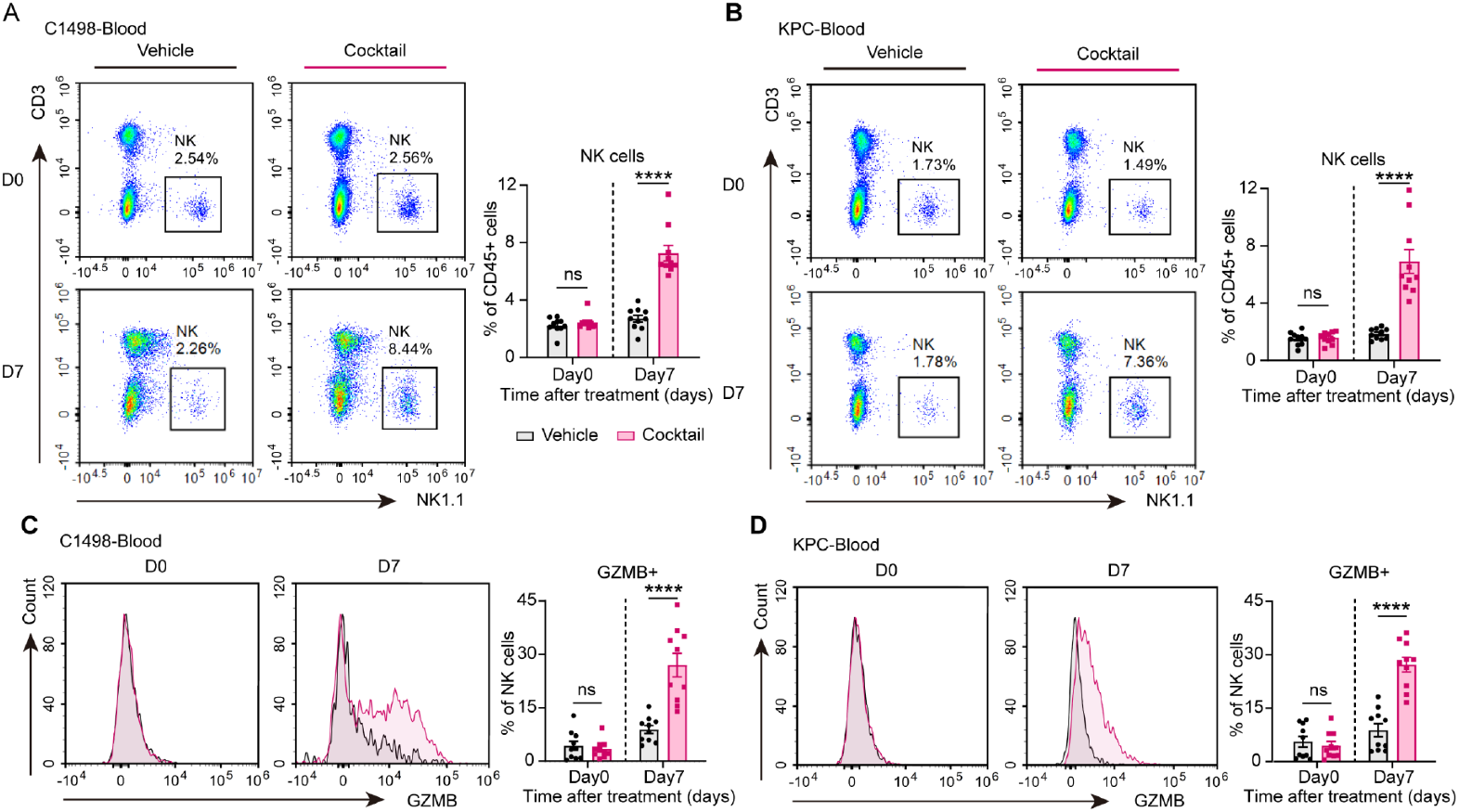
Topical cocktail-based in vivo vaccine expands functional NK cells in blood in mouse models. **A. B**. Representative flow cytometry graphs and quantification of percentages of NK cells in CD45^+^ cells in blood of C1498 (**A**) and KPC (**B**) xenograft C57 mice on day0 and 7 after vehicle or vaccine treatment (n=10). **C. D**. Representative flow cytometry graphs and quantification of percentages of GZMB^+^ cells in NK cells in blood of C1498 (**C**) and KPC (**D**) xenograft C57 mice on day0 and 7 after vehicle or vaccine treatment (n=10). Gating strategy is depicted in **Fig. S2F**. All error bars represent means ± s.e.m. Statistical significance was determined by the two-tailed t-test. **** p<0.0001, ns means no significance.

Notably, functional GZMB^+^NK cells were significantly augmented in both vaccine-treated mouse AML and PDAC models (Fig. 4C, D). In mouse AML model, the frequency of GZMB^+^NK cells increased by 3.04-fold following vaccine treatment (Fig. 4C) (8.88% GZMB^+^NK cells (vehicle) vs. 27.00% GZMB^+^NK cell (vaccine), P < 0.0001). Similarly, the frequency of GZMB^+^NK cells increased by 3.09-fold following vaccine treatment (Fig. 4D) (8.82% GZMB^+^NK cell (vehicle) vs. 27.22% GZMB^+^NK cell (vaccine), P < 0.0001) in mouse PDAC model.

To confirm whether the in vivo vaccine affects NK cells in spleens, we analyzed NK cells and GZMB^+^NK cells in spleen samples from vaccine-treated and control mice. Consistently, the overall frequency of NK cells increased in spleens in both vaccine-treated mouse AML and PDAC models (S. Fig. 1A, B). In mouse AML model, the frequency of NK cells increased by 1.57-fold following vaccine treatment (S. Fig. 1A) (1.97% NK cell (vehicle) vs. 3.09% NK cell (vaccine), P < 0.05). The frequency of NK cells increased by 1.46-fold following vaccine treatment (S. Fig. 1B) (1.59% NK cell (vehicle) vs. 2.32% NK cell (vaccine), P < 0.05) in mouse PDAC model. Importantly, the frequency of GZMB^+^NK cells increased by 3.55-fold following vaccine treatment (S. Fig. 1C) (6.35% GZMB^+^NK cell (vehicle) vs. 22.55% GZMB^+^NK cell (vaccine), P < 0.05) in mouse AML model. Similarly, the frequency of GZMB^+^NK cells increased by 4.18-fold following vaccine treatment (S. Fig. 1D, E) (5.62% GZMB^+^NK cell(vehicle) vs. 23.49% GZMB^+^NK cell (vaccine), P < 0.001) in mouse PDAC model (S. Fig. 1D).

We next determined NK cells and GZMB^+^NK cells in bone marrows from vaccine-treated and control mice using flow cytometry. In mouse AML model, the frequency of NK cells increased by 1.29-fold following vaccine treatment (S. Fig. 1E) (0.77% NK cell (vehicle) vs. 0.99% NK cell (vaccine), P > 0.05). In mouse PDAC models, the frequency of NK cells decreased by 0.90-fold following vaccine treatment (0.88% NK cell (vehicle) vs. 0.80% NK cell (vaccine), P > 0.05) (S. Fig. 1F). The frequency of GZMB^+^NK cells increased by 6.11-fold following vaccine treatment (3.43% GZMB^+^NK cell (vehicle) vs. 20.96% GZMB^+^NK cell (vaccine), P < 0.05) in mouse AML model (S. Fig. 1G). Likely, the frequency of GZMB^+^NK cells increased by 4.37-fold following vaccine treatment (7.21% GZMB^+^NK cell(vehicle) vs. 31.54% GZMB^+^NK cell (vaccine), P < 0.0001) (S. Fig. 1H).

Collectively, these results reveal that the in vivo vaccine can systemically activate and expand functional NK cells in blood, spleen and bone marrow in mouse AML and PDAC models.

### The in vivo vaccine boosts cytotoxic CD8+ T cells systemically in tumor-bearing mice

Cytotoxic CD8^+^ T cells play a central role in cancer immunotherapy^35, 36, 37^. To explore whether the in vivo vaccine-induced anti-tumor activity is involved in CD8^+^ T cells, we next investigated the frequency of IFN-γ^+^ CD8^+^ and GZMB^+^CD8^+^ T cells in the bloods between vehicle- and vaccine-treated mouse AML and PDAC models after 7 days of treatment with flow cytometry. No differences in the frequencies of CD3^+^ T cells were detected between vaccine-treated and vehicle-treated groups in both mouse AML and PDAC mice (S. Fig. 2A, b). In contrast, the overall frequency of CD45^+^CD8^+^ T cells significantly increased in the bloods in both vaccine-treated mouse AML and PDAC models (Fig. 5A, B). In mouse AML model, the frequency of CD45^+^ CD8^+^ T cells increased by 1.23-fold following vaccine treatment (11.34% CD45^+^ CD8^+^ T cells (vehicle) vs. 13.91% CD45^+^ CD8^+^ T cells (vaccine), P < 0.05) (Fig. 5A). Similarly, the frequency of CD45^+^ CD8^+^ T cells increased by 1.27-fold following vaccine treatment (10.29% CD45^+^ CD8^+^ T cells (vehicle) vs. 13.10% CD45^+^ CD8^+^ T cells (vaccine), P < 0.01) in mouse PDAC model (Fig. 5B).

**Figure 5.**
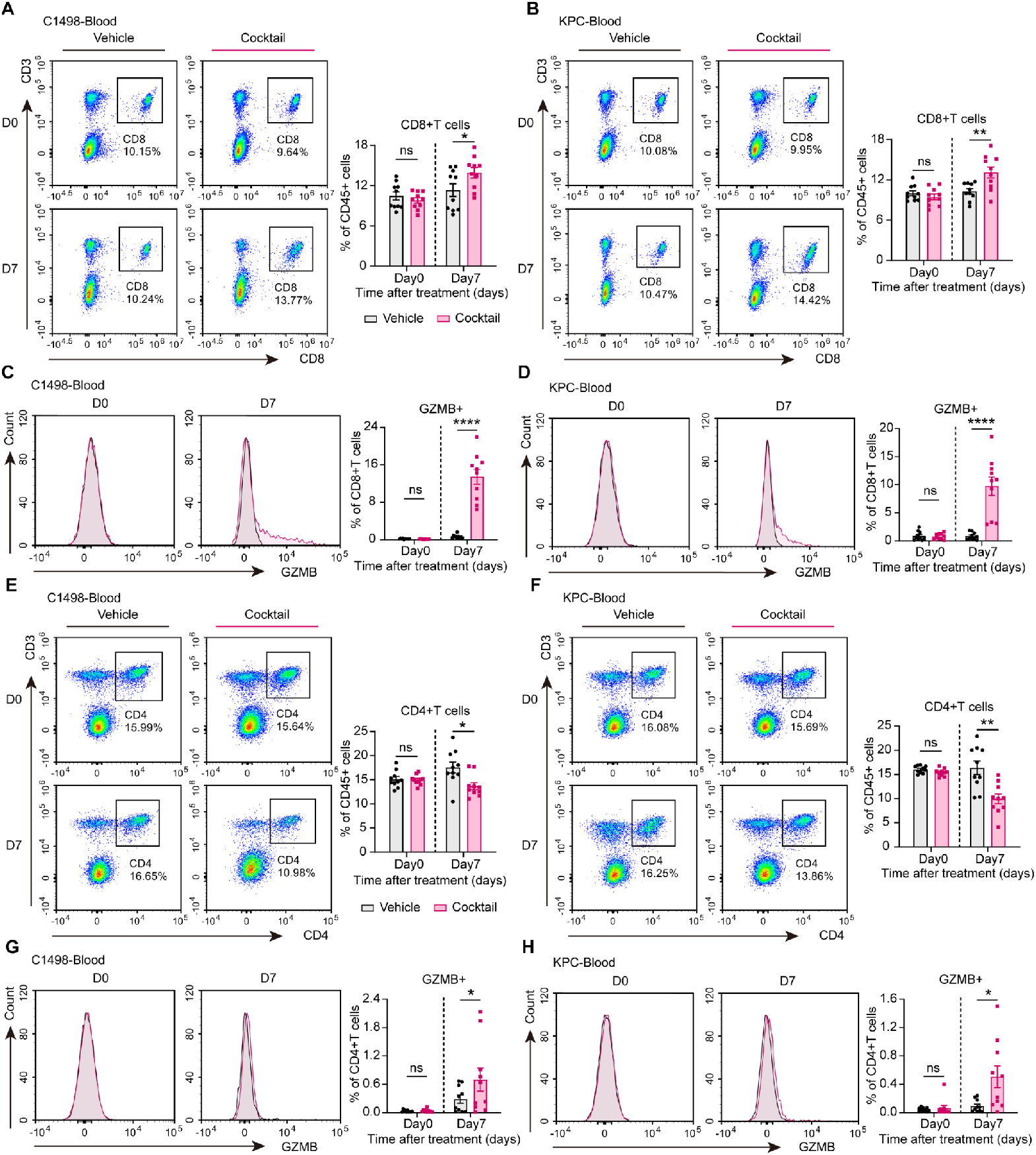
Topical cocktail expands functional CD8+ and CD4+CTL cells in blood in AML and PDAC mouse models. **A. B**. Representative flow cytometry graphs and quantification of percentages of CD8^+^T cells in CD45^+^ cells in blood of C1498 (**A**) and KPC (**B**) xenograft C57 mice on day0 and 7 after vehicle or vaccine treatment (n=10). **C. D**. Representative flow cytometry graphs and quantification of percentages of GZMB^+^ cells in CD8^+^T cells in blood of C1498 (**C**) and KPC (**D**) xenograft C57 mice on day0 and 7 after vehicle or vaccine treatment (n=10). **E. F**. Representative flow cytometry graphs and quantification of percentages of CD4^+^T cells in CD45^+^ cells in blood of C1498 (**E**) and KPC (F) xenograft C57 mice on day0 and 7 after vehicle or vaccine treatment (n=10). **G. H**. Representative flow cytometry graphs and quantification of percentages of GZMB^+^ cells in CD4^+^T cells in blood of C1498 (**G**) and KPC (**H**) xenograft C57 mice on day0 and 7 after vehicle or vaccine treatment (n=10). Gating strategy is depicted in **Fig. S2F**. All error bars represent means ± s.e.m. Statistical significance was determined by the two-tailed t-test, * p<0.05, ** p<0.01, **** p<0.0001, ns means no significance.

We next determined the frequency of functional IFN-γ^+^ CD8^+^ T cells and GZMB^+^ CD8^+^ T cells. To our surprise, functional GZMB^+^ CD8^+^ T cells markedly increased in both vaccine-treated mouse AML and PDAC models (Fig. 5C, D), but no differences in the frequencies of IFN-γ^+^CD8^+^ T cells were observed between vaccine-treated and vehicle-treated groups in tumor-bearing mice (S. Fig. 2C, D). In mouse AML model, the frequency of GZMB^+^ CD8^+^ T cells increased by 19.24-fold following vaccine treatment (Fig. 5C) (0.70% GZMB^+^ CD8^+^ T cells (vehicle) vs. 13.47% GZMB^+^ CD8^+^ T cells (vaccine), P < 0.0001). Consistently, the frequency of GZMB^+^ CD8^+^ T cells increased by 11.61-fold following vaccine treatment (0.84% GZMB^+^ CD8^+^ T cells (vehicle) vs. 9.75% GZMB^+^ CD8^+^ T cells (vaccine), P < 0.0001) in mouse PDAC model (Fig. 5D).

We next examined frequencies of CD8^+^ T cells and GZMB^+^ CD8^+^ T cells in spleens from vaccine-treated and control mice using flow cytometry. The overall frequency of CD8^+^ T cells were not significantly changed between vaccine-treated and vehicle-treated spleens in both mouse AML and PDAC models (S. Fig. 3A, B). However, the frequency of GZMB^+^ CD8^+^ T cells significantly increased in vaccine-treated spleens in both mouse AML and PDAC models (S. Fig. 3C, D). The frequency of GZMB^+^CD8^+^ T cells increased by 3.13-fold following vaccine treatment (S. Fig. 3C) (0.32% GZMB^+^ CD8 T cells (vehicle) vs. 1.00% GZMB^+^ CD8^+^ T cells (vaccine), P < 0.01) in mouse AML model. The frequency of GZMB^+^ CD8^+^ T cells increased by 7.15-fold following vaccine treatment (S. Fig. 3D) (0.20% GZMB^+^ CD8^+^ T cells(vehicle) vs. 1.43% GZMB^+^ CD8^+^ T cells (vaccine), P < 0.01) in mouse PDAC model.

We further analyzed frequencies of CD8^+^ T cells and GZMB^+^ CD8^+^ T cells in bone marrows from vaccine-treated and control mice using flow cytometry. The overall frequency of CD8^+^ T cells increased in mouse AML model (S. Fig. 3E), but slightly decreased in mouse PDAC model between vaccine-treated and vehicle-treated bone marrows (S. Fig. 3F). However, the frequency of GZMB^+^ CD8^+^ T cells significantly increased in vaccine-treated BM in both mouse AML and PDAC models (S. Fig. 3G). The frequency of GZMB^+^CD8^+^ T cells increased by 4.86-fold following vaccine treatment (S. Fig. 3G) (1.44% GZMB^+^ CD8^+^ T cells (vehicle) vs. 7.00% GZMB^+^ CD8^+^ T cells (vaccine), P < 0.01) in mouse AML model. The frequency of GZMB^+^ CD8^+^ T cells increased by 12.79-fold following vaccine treatment (S. Fig. 3H) (0.97% GZMB^+^ CD8^+^ T cells (vehicle) vs. 12.41% GZMB^+^ CD8^+^ T cells (vaccine), P < 0.0001) in mouse PDAC model.

### The in vivo vaccine elicits cytotoxic CD4+ T cells systemically in tumor-bearing mice

Given that cytotoxic CD4^+^ T cells can directly kill CD8^+^ T cell-resistant cancer cells ^38, 39, 40^, we next investigated the frequency of CD4^+^ T cells and cytotoxic GZMB^+^ CD4^+^ T cells in the bloods between vehicle- and vaccine-treated mouse AML and PDAC models after 7 days of treatment. The overall frequency of CD45^+^ CD4^+^ T cells significantly decreased in the bloods in both vaccine-treated mouse AML and PDAC models (Fig. 5E, F). In mouse AML model, the frequency of CD45^+^ CD4^+^ T cells decreased by 0.78-fold following vaccine treatment (16.39 % CD45^+^ CD4^+^ T cells(vehicle) vs. 9.98% CD45^+^ CD4^+^ T cells (vaccine), P < 0.05) (Fig. 5E).

Similarly, the frequency of CD45^+^ CD4^+^ T cells decreased by 0.61-fold following vaccine treatment (17.53% CD45^+^ CD4^+^ T cells(vehicle) vs. 13.67% CD45^+^ CD4^+^ T cells (vaccine), P < 0.01) in mouse PDAC model (Fig. 5F). Surprisingly, cytotoxic GZMB^+^ CD4^+^ T cells were significantly increased in both vaccine-treated mouse AML and PDAC models (Fig. 5G, H). The frequency of GZMB^+^ CD4^+^ T cells increased by 2.41-fold following vaccine treatment (Fig. 5G) (0.29% GZMB^+^ CD4^+^ T cells(vehicle) vs. 0.70% GZMB^+^ CD4^+^ T cells (vaccine), P < 0.05) in mouse AML model. The frequency of GZMB^+^ CD4^+^ T cells increased by 5.67-fold following vaccine treatment (Fig. 5H) (0.09% GZMB^+^ CD4^+^ T cells(vehicle) vs. 0.51% GZMB^+^ CD4^+^ T cells (vaccine), P < 0.05) in mouse PDAC model.

Similar decrease of CD4^+^ T cells were observed in spleens and bone marrow in both mouse AML and PDAC models (S. Fig. 4A, B, E, F). However, the frequency of cytotoxic GZMB^+^ CD4^+^ T cells in spleens increased by 3.20-fold following vaccine treatment (S. Fig. 4C) (0.05% GZMB^+^ CD4^+^ T cells(vehicle) vs. 0.16% GZMB^+^ CD4^+^ T cells (vaccine), P < 0.01) in mouse AML model. The frequency of GZMB^+^ CD4^+^ T cells in spleen increased by 4.67-fold following vaccine treatment (S. Fig. 4D) (0.03% GZMB^+^CD4^+^ T cells (vehicle) vs. 0.14% GZMB^+^ CD4^+^ T cells (vaccine), P < 0.05) in mouse PDAC model.

The frequency of cytotoxic GZMB^+^ CD4^+^ T cells in bone marrow increased by 1.93-fold following vaccine treatment (S. Fig. 4G) (0.42% GZMB^+^ CD4^+^ T cells(vehicle) vs. 0.81% GZMB^+^CD4^+^ T cells (vaccine), P < 0.05) in mouse AML model. The frequency of GZMB^+^CD4^+^ T cells in spleen increased by 2.52-fold following vaccine treatment (S. Fig. 4H) (0.27% GZMB^+^ CD4^+^ T cells(vehicle) vs. 0.68% GZMB^+^ CD4^+^ T cells (vaccine), P < 0.01) in mouse PDAC model.

Together, these data indicate that the in vivo vaccine can elicit cytotoxic CD4^+^ T cells systemically.

### The in vivo vaccine expands DC cells and reduces Treg T cells in mouse AML and PDAC models

Dendritic cells (DCs) are potent antigen-presenting cells and play a pivotal role in the cancer-immunity cycle by priming and activating antigen-specific effector T cells, and their deficiency in the tumor microenvironment is a key reason for immune escape. Restoring DC immunity could improve immunotherapy efficacy ^41, 42, 43, 44, 45^. To determine whether the in vivo vaccine influences DCs, we next investigated the frequency of circulating DC cells between vehicle- and vaccine-treated mouse AML and PDAC models at day 7 after treatment with vaccine in the bloods. We used flow cytometry to quantify DCs (CD11c^+^ cells). In AML mouse model, the frequency of DCs increased by 3.00-fold at day 7 (0.11% DC (vehicle) vs. 0.33% DC (vaccine), P < 0.01) (S. Fig. 5A). Similarly, the frequency of DC increased by 2.75-fold at day 7 (0.08 % DC (vehicle) vs. 0.22% DC (vaccine), P < 0.01) in mouse PDAC model (S. Fig. 5B). However, frequency of DC was not significantly changed between vaccine-treated and vehicle-treated spleens and bone marrow in both mouse AML and PDAC model (S. Fig. 5C, D, E, F).

Treg cells have been shown to be the major driver of immune escape in cancer patients receiving immunotherapy; we next assessed the effects of vaccine on these inhibitory T cells in spleen and bone marrow, which are TEM of leukemia, in mouse AML model. Treg cells significantly decreased from 1.55% (vehicle) to 1.07% (vaccine) (P < 0.05) (S. Fig. 5G) in spleens. Consistent with the spleen findings, Treg cells were also decreased from 0.15% (vehicle) to 0.07% (vaccine) (P < 0.05) (S. Fig. 5H) in bone marrows in mouse AML model. These findings indicate that in vivo vaccine can reduce inhibitory T cells in tumor microenvironments.

### The in vivo vaccine uniquely generates a super immune cell network in TME and DLNs of tumor-bearing mice

We next studied phenotypes of immune cells by analyzing histopathological and immunological changes of tumor tissues between vaccine-treated mice and vehicle-treated mice in mouse PDAC models using multiplex immunofluorescence (mIF). The tumors were collected and processed for H&E staining and immunofluorescence staining at day 7 after vaccine treatment. We observed large areas of necrosis in the central region of tumor tissue but not in vehicle-treated tumors (Fig. 6A-H). mIF staining results showed that various immune cells, including CD8^+^ T cells, CD4^+^ T cells, NK cells, CD11c^+^ DCs and were significantly enriched within vaccine-treated tumor tissues as compared the vehicle-treated tumors (Fig. 6A). The in vivo vaccine treatment increased CD8^+^T, GZMB^+^ CD8 T, CD4^+^ T, GZMB^+^ CD4 T, NK, GZMB^+^ NK, and DC cell populations by 21.67-, 4.37-, 14.74-, 6.74-, 6.92-, 4.71- and 35.37-folds in TME, respectively, respectively, in tumor tissues, as compared with vehicle-treated tumor tissues (Fig. 6B-H).

**Figure 6.**
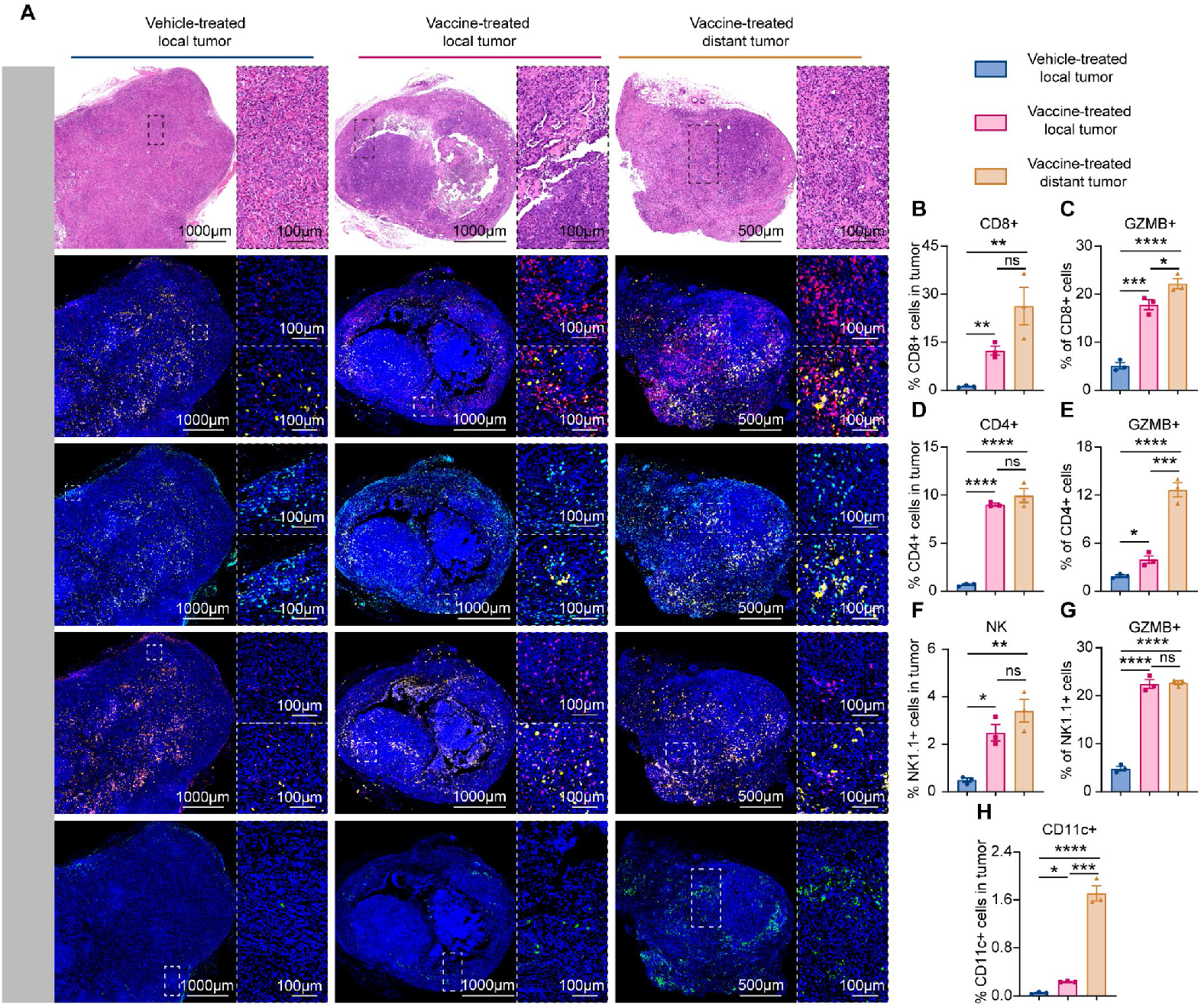
The in vivo vaccine generates a super pan-immune cell network in tumor microenvironments. **A**. Representative and the corresponding local enlarged images of H&E, GZMB^+^ (yellow) CD8^+^ (Red) T cells, GZMB^+^ (yellow) CD4^+^ (Cyan) T cells, GZMB^+^ (yellow) NK (Magenta) cells and CD11c (Green) staining of mice tumor from each treatment group, counterstained with DAPI (Blue). **B-H**. Quantification of CD8^+^ (**B**), GZMB^+^ CD8^+^ (**C**), CD4^+^ (**D**), GZMB^+^ CD4^+^ (**E**), NK (**F**), GZMB^+^ NK (**G**), CD11c^+^ (**H**) cells in tumor tissue (n=3). All error bars represent means ± s.e.m. Statistical significance was determined by one-way ANOVA. * p<0.05, ** p<0.01, *** p<0.001, **** p<0.0001, ns means no significance.

To further confirm above observations, we analyzed histological and immunological changes of implanted tumors in vaccine-cured mice after 7 days of engraftment and tumors in vehicle-treated naïve mice in mouse PDAC models. Consistent with above observations, large areas of necrosis in the central region were observed in implanted tumors in vaccine-cured mice but not in vehicle-treated tumors in naïve mice. mIF staining results showed that various immune cells were enriched at the edges of vaccine-treated tumor tissues, but not in the vehicle-treated tumors (S. Fig. 6A, B). The in vivo vaccine treatment increased GZMB^+^ NK, DC, GZMB^+^ CD8 and GZMB^+^ CD4 cell levels by 1.93, 10.22, 1.44 and 1.24-folds, respectively, in tumor tissues, as compared with vehicle-treated tumor tissues (S. Fig. 6C-F).

We next compared immunological changes of draining lymph nodes between vaccine-treated mice and vehicle-treated mice in mouse PDAC models using mIF. The lymph nodes were collected and processed for H&E staining and immunofluorescence staining at day 7 after vaccine treatment. No significant histological changes were observed in DLNs between vaccine-treated mice and vehicle-treated mice (S. Fig. 7A). However, mIF staining results showed that various immune cells were enriched in vaccine-treated DLNs, but not in the vehicle-treated DLNs (S. Fig. 7A). The in vivo vaccine treatment increased GZMB^+^NK, DC, GZMB^+^CD8 and GZMB^+^CD4 cell levels by 2.28, 2.05, 1.87 and 3.79-folds, respectively, in vaccine-treated DLNs, as compared with vehicle-treated DLNs (S. Fig. 7B-E).

These findings, together with the facts that GZMB^+^ NK, DC, GZMB^+^ CD8^+^ and GZMB^+^ CD4^+^ T cells increased in the blood and spleen in vaccine-treated tumor-bearing mice, indicate that the in vivo vaccine has unique ability to generate a super immune cell network in both TME and DLNs.

### Cocktail acts directly on human innate and adaptive immune cells to enhance anti-tumor response

To test whether the tumor cell killing observed in mouse AML and PDAC models would translate to human patients, we next investigated the effects of cocktail on the tumor lysis ability of PBMCs via co-culturing PBMCs with PDAC MIA-paca2 cells (E:T ratio of 5:1). We isolated PBMCs from PDAC patients as well as healthy donors. After PBMCs were treated with cocktail or single component for indicated timepoint, we measured anti-tumor activity of PBMCs against PDAC cells and AML cells. We found that both cocktail and single component significantly improved cytotoxicity of PBMCs against PDAC cells. Anti-tumor activity of PBMCs increased by 1.15-fold, 1.18-fold and 1.50-fold, respectively, after treatment with cocktail, PT100 (at 0.5μM) and RMQ (at 0.5μM) (S. Fig. 8A, B). Similar results were also observed in PBMCs healthy donors. Anti-tumor activity of PBMCs increased by 1.40-fold, 1.10-fold and 1.56-fold, respectively, after treatment with cocktail, PT100 (at 0.5μM) and RMQ (at 0.5μM) (S. Fig. 8A, B). These results suggest that the cocktail-based in vivo formulation can improve immune cell anti-tumor activity in blood from both cancer patients and normal individuals.

We next investigated whether component of cocktail could affect the tumor lysis ability of NK cells via co-culturing NK92 cells with PDAC MIA-paca2 cells. We treated human NK92 cells with RMQ or PT100 for 24 hours and then investigated tumor lysis ability. We found that both RMQ and PT100 markedly increased NK92 cell anti-tumor activity against PDAC MIA-paca2. Anti-tumor activity of NK92 cells increased by 1.23-fold and 2.35-fold, respectively, after treatment with PT100 (at 0.5μM) and RMQ (at 0.5μM) (S. Fig. 9A, B). However, both RMQ and PT100 showed no obvious cytotoxicity against cancer cells at the test concentrations (S. Fig. 9 A, B). These findings suggest that agonist PT100 and RMQ can directly enhance NK cell anti-tumor function.

To evaluate whether PT100 and RMQ affect T cell anti-tumor activity, we next investigated whether component of cocktail could affect the tumor lysis ability of CD3^+^ T cells via co-culturing CD3^+^ T cells with PDAC MIA-paca2 cells. We treated primary CD3^+^ T cells from patients with PDAC patients with RMQ or PT100 for 24 hours and then investigated tumor lysis ability. We found that both RMQ and PT100 markedly increased CD3^+^T cell anti-tumor activity against PDAC MIA-paca2. Anti-tumor activity of CD3^+^T cells increased by 1.57-fold and 1.69-fold, respectively, after treatment with PT100 (at 0.5μM) and RMQ (at 0.5μM) (S. Fig. 10 A, B). These results indicate that agonist PT100 and RMQ can directly enhance CD3^+^T cell anti-tumor function.

### Topical cocktail-based in vivo vaccine is well tolerated in mice

To assess the tolerability of transdermal cocktail, mouse toxicity was evaluated by topical application for once a week for 3 weeks and by monitoring for lethality, mouse weight, blood counts, blood liver enzyme (ALT, AST), serum creatinine (Scr) and myocardial enzyme spectrum (CK, CK-MB, LDH). No systemic toxicity was observed tested mouse models.

Analyses of body weights, leukocyte subsets, blood liver enzymes ALT (41.1U/L (vehicle) vs. 35.3U/L (vaccine)), AST (106.8U/L (vehicle) vs. 99.1U/L (vaccine)), Scr (21.0umol/L (vehicle) vs. 19.9umol/L (vaccine)) and myocardial enzyme spectrum (CK (291.3U/L (vehicle) vs. 308.8U/L (vaccine)), CK-MB (306.8U/L (vehicle) vs. 322.2U/L (vaccine)), LDH (628.2U/L (vehicle) vs. 599.2U/L (vaccine))) did not reveal any substantial differences between the cocktail-treated group and control group (S. Fig. 11A-D), suggesting negligible systemic toxicity of the cocktail. Notably, all of the mice dosed with cocktail even at as high as twice dose of cocktail survived well except for a transient weight loss of 3.35% compared to control animals at 7 days after receiving the first dose of topical cocktail, however, the weight loss was fully reversible after completing the treatment cycle (S. Fig. 11E). Moreover, no adverse effects were observed at treated skin sites with cocktail. These results suggest that transdermal cocktail-based in vivo vaccine is safe in mouse models.

Current therapeutic cancer vaccines have been challenged by the limited therapeutic efficacy due to the deficiency of the cancer-immunity cycle in most tumors, therefore, effective immunotherapy requires the activation of the entire cancer-immunity cycle, including the release of tumor antigens from cancer cells, antigen processing and presentation by antigen-presenting cells (APCs), the generation of tumor-specific immune responses and tumor lysis, which increases tumor antigen release and amplifies the cycle. Direct in vivo cancer vaccination is the best strategy to restore the entire cancer-immunity cycle. The in vivo vaccine works by inducing tumor cells to release their immunogenic antigens in a way that promotes dendritic cells and other APCs to present these antigens to T cells and other immune cells, leading to activation and expansion of immune cells. This co-localization of tumor antigens and APCs can be finished directly at the tumor site.

Therefore, we designed the novel in vivo vaccine with a topical small molecule cocktail as antigen-agnostic vaccine to boost the entire cancer-immunity cycle. We designed NK and DC activators RMQ and PT100 to trigger immunogenic cell death (ICD) and release tumor antigens, and promote process and present tumor antigens, which systemically and extensively activate innate and adaptive immunity to clear tumors cross the body. In addition, we designed DTH as a strong inflammatory inducer to improve innate and adaptive immune responses with immune memory. As expected, a systemic activation of diverse innate and adaptive anti-tumor immunity was observed in tumor-bearing AML and PDAC mouse models. These findings suggest that the in vivo vaccine robustly reprogram the immune microenvironment of entire body to develop high-magnitude, functional immune cell responses.

Of note, our studies reveal that the in vivo vaccine treatment induces marked increases of DCs in tumor microenvironment (TEM), DLN, blood, spleen and bone marrow, indicating that the vaccine systemically activates DC cells. This increase greatly improves tumor-specific T cell priming and subsequent T cell supply to the TME.

Another interesting aspect of this study is the observation of an extensive activation of diverse innate immune cell populations such as NK cells, CD11b^+^/Ly6G^+^/Ly6C^+^ myeloid inflammatory cell, which are crucial to clear circulating cancer cells. In addition, we also demonstrate that the in vivo vaccine elicits cytotoxic CD4^+^ T cells systemically, which can directly kill cancer cells resistant to CD8^+^ T cells.

Despite the potent immune responses induced, no vaccine-related side effects, such as cytokine release syndrome, were observed. Furthermore, no autoimmune-related safety events were observed. Finally, the cocktail-based in vivo vaccine is well simplified clinical medication for cancer patients.

In summary, we developed a novel in vivo vaccine with a topical triple-small molecule cocktail formulation as antigen-agnostic cancer vaccine, which can rapidly eliminate hematological and solid tumors in preclinical mouse models via resetting the entire cancer-immunity cycle. The novel in vivo vaccine might represent a promising avenue for pan-cancer therapy, and will provide the framework for further clinical trial.

## Materials and Methods

### Cell lines and culture

The murine leukemia cell line (C1498) and murine pancreatic cancer (KPC) were maintained in DMEM medium (11965092, Gibco, USA) with 10% FBS (BX0500C, Braserum), 1% penicillin/streptomycin (PS2004HY, TBD Science). Human pancreatic cancer (MIA-paca2) was maintained in DMEM medium with 10% FBS, 1% penicillin/streptomycin. NK92-MI cell line was kept in MEM-α medium (12561056, Gibco, USA), supplemented with supplemented with 12.5% FBS, 12.5% HS (26050088, Gibco) and 1% penicillin/streptomycin. All cells were cultured in a humidified atmosphere containing 5% CO_2_ at 37 °C and routinely tested for mycoplasma.

### PBMCs and CD3+ T Cell Isolation and Culture

Peripheral blood mononuclear cells (PBMCs) were isolated and purified from blood samples collected from healthy donors and PDAC patients by Ficoll-Paque density-gradient centrifugation (LTS1077, TBD Science), cultured in Iscove-modified Dulbecco medium (IMDM) (CR-12200, Cienry) containing 15%FBS. CD3^+^ T cells were collected from PBMC using CD3-specific microbeads (L00896, GenScript) and cultured in IMDM with 20% FBS (10091148, Gibco), 1% penicillin/streptomycin, IL-2 (300U mL−1, QUANGANG PHARMACEUTICAL). CD3/CD28 Dynabeads (11161D, Invitrogen) were added for the activation and expansion. Cell purity was determined > 95% by flow cytometry.

### Preparation of topical small molecule cocktail formulation

The topical small molecule cocktail is three-component pan-immunity activators comprising resiquimod (RMQ) (Cas: 144875-48-9, Aladdin) and PT100 (Cas: 150080-09-4, Aladdin) as well as DTH (Cas: 1143-38-0, Aladdin) as co-adjuvant for non-invasive dermal vaccination. These components were formulated into one with Vaseline cream (A510146-0500, Sangon Biotech), which can be easily used for patients.

### Non-invasive topical administration of in vivo vaccine

Briefly, dorsal hair was removed with electric clippers. Mice were treated with 0.1ml of cocktail cream or vehicle cream, which applied onto skin covering a surface area of up to 1cm^2^. The amount of each component (RMQ, PT100 and DTH) in 0.1ml cocktail cream is far below the amount commonly used in animal models and in the treatment of psoriatic patients.

### Establishment of mouse model

Male and female C57BL/6 mice at 6–8 weeks were obtained from Shanghai SLAC Laboratory Animal Co., Ltd. (Shanghai, China). KPC cells harboring mutant KRAS^G12D^ and TP53^R172H^ were overexpressed with pCDH-luciferase plasmids to develop KPC-Luci cells. Luciferase-tagged KPC cells (KPC-Luc, 2×10^6^ per mouse) were implanted subcutaneously in the left flank of 6-week-old C57BL mice. C1498 cell line was labeled with pCDH-luciferase-GFP plasmids and luciferase-GFP-tagged C1498 cells (C1498-Luc, 5×10^6^ per mouse) injected through the tail vein into mice. After obvious tumor signals could be detected (day 7 after cells injection), mice were randomly divided into different groups to receive therapy. Tumor burden was monitored by in vivo fluorescence imaging using IVIS lumina LT series III in vivo imaging system (Perkin 710 Elmer, USA), mice weight and activity were recorded at different time points. The tumor width and length were measured by one experienced animal technician. Tumor volume was calculated using the formula: (length × width2)/2. The venous blood from the orbital vein were collected on days 0, 7 and 14 for flow cytometry analyses, leukocyte subsets, blood liver enzymes ALT and AST, serum creatinine and myocardial enzyme spectrum (CK, CK-MB, LDH). The tumor, spleen, liver, Bone narrow, lymph nodes were obtained on day14 after euthanized for H&E staining, and multiplex immunofluorescence (mIF). At the end of the experiment, mice were given euthanasia.

### TUNEL assay

The terminal Deoxynucleotidyl Transferase-Mediated dUTP Nick End Labeling (TUNEL) staining was carried out on mouse tumors using the TUNEL Bright Green Apoptosis Detection Kit (A112-03, Vazyme), according to the manufacturer’s instructions. Briefly, slides containing paraffin-embedded tissues were dewaxed in deparaffinizing agent and placed in the citrate antigen retrieval solution for 1h. After washing three times in TBST, the slides were incubated with TUNEL reaction mixture at 4°C overnight. Then slides were incubated with DAPI for 5min after washing three times in TBST, and imaged with a fluorescence microscope (BX53, Olympus).

### Immunohistochemistry and multiplex immunofluorescence staining

Mouse tumors and lymph nodes were fixed in 4% PBS-buffered formalin (E672002, Sangon Biotech), embedded in paraffin, sectioned and stained with hematoxylin and eosin (H&E) (BP0211, Biossci). For mIF, sections were dewaxed and washed three times in distilled water. Antigen retrieval was performed in EDTA Buffer (pH9, Biossic) using a pressure cooker for 1min. After washed three times in TBST, Sections were incubated in 3% hydrogen peroxide for 10min and blocked in 10% sheep serum for 30min. Primary antibodies were diluted in TBST and incubated at 4°C overnight. After washed three times in TBST, Sections were incubated in Goat Anti-Rabbit IgG H&L HRP secondary antibodies (1:4000, Abcam, RRID: AB_2819160) for 45min and then incubated in opal working solution which containing tyramide signal amplification substrates for 10min at room temperature. The slides were then antigen retrieved and the process was repeated up. At the end of the experiment, samples were stained with DAPI and visualized on a Zeiss confocal microscope. The sample spectral unmixing and quantification of signals were conducted with 3D HISTECH (Hungary), using the ImageJ software.

The following primary antibodies were used: CD11b (1:8000, Abcam, RRID: AB_2650514), Ly6G (1:5000, Abcam, RRID: AB_2923218), Ly6C (1:100, Abcam, ab314120), CD11c (1:1000, CST, 97585S), CD4 (1:4000, Abcam, RRID: AB_2686917), CD8 (1:4000, Abcam, RRID: AB_2860566), NK1.1 (1:200, Abcam, RRID: AB_3094493), Granzyme B (1: 4000, Abcam, RRID: AB_2860567).

### Flow cytometry analyses

Flow cytometry was performed and analyzed using NovoCyte cytometer (ACEA, Agilent, Santa Clara, CA, USA). The following antibodies were purchased from Biolegend (San Diego, CA, USA): mouse CD16/32 (clone S17011E, RRID: AB_2783137), Zombie Aqua™ Fixable Viability Kit (423101), mouse CD45-Pacific Blue (clone 30F11, RRID: AB_493535), mouse CD3-FITC (clone 17A2, RRID: AB_312660), mouse CD4-PerCP/Cyanine5.5 (clone GK1.5, RRID: AB_893324), mouse CD8a-PE (clone 53-6.7, RRID: AB_312747), mouse CD8a-APC/Fire 750 (clone 53-6.7, RRID: AB_2572113), mouse NK1.1-PE/Cyanine7 (clone PK136, RRID: AB_389364), mouse CD25-APC (clone PC61, RRID: AB_312860), mouse FOXP3-PE (clone MF-14, RRID: AB_1089118), human/mouse Granzyme B-APC (clone QA16A02, RRID: AB_2687028), mouse IFN-γ-APC/Cyanine7 (clone XMG1.2, RRID: AB_2616698), mouse/human CD11b-FITC (clone M1/70, RRID: AB_312789), mouse Ly6G-Brilliant Violet 785 (clone 1A8, RRID: AB_2566317), mouse Ly6C-APC/Cyanine7 (clone HK1.4, RRID: AB_10643867), mouse CD11c-Brilliant Violet 570 (clone N418, RRID: AB_10900261), mouse F4/80-PE (clone BM8, RRID: AB_893498), mouse I-A/I-E-APC (clone M5/114.15.2, RRID: AB_313328), human CD45-Pacific Blue (clone HI30, RRID: AB_2561357), human CD3-APC (clone OKT3, RRID: AB_1937213).

### Cell Proliferation and Viability Assays by MTT

RMQ and PT100 were dissolved in DMSO to a final concentration of 10 mg mL–1 and stored at −20 °C. For MTT (3-(4,5-dimethylthiazol-2-yl)-2,5-diphenyltetrazolium bromide) assay, cells in an exponential phase of growth were seeded in a 96-well plate with 200 μL medium and cultured for 72 h in the presence of various concentrations of drugs. Then, cells were incubation with 0.5 mg/mL MTT (A600799, Sangon Biotech) for 4 h at 37 °C followed by incubation with lysis buffer (10% SDS, 5% Isobutanol, 0.012M HCl) for 16 h and absorbance was measured at 560 nm. The IC50 was calculated using GraphPad Prism version 8.

### Cytotoxicity assays

Luciferase-expressing MIA-paca2 (1×10^4^) were co-cultured with effector cells and drugs at various E:T ratios in a 96-well plate. The maximal cell killing group (Max) was treated with Triton X-100, while no effector cells were added for the minimal cell killing group (Min). At the end point, 100 μL 1.0 mg/mL of D-Luciferin Salt Bioluminescent substrate (PerkinElmer) was added after the supernatant was discarded, and measured by a Specctramax i3 instrument (Molecular Devices, Sunnyvale, USA). The specific lysis was calculated using the following formulas: specific lysis = (Min − test)/(Min − Max) × 100%.

### Crystal Violet Staining

GFP-expressing MIA-paca2 (1×10^4^) were co-cultured with effector cells and drugs at various E:T ratios in a 96-well plate. Cells were then imaged with Zeiss Confocal Laser Scanning Microscope 710 (LSM710, Germany). ZEISS ZEN Microscope software was used for acquisition and analysis. The supernatant was carefully discarded and 100μL methanol was added for 10min at room temperature. Then removed the methanol, covered with 1% crystal violet staining solution (C0121, Beyotime) and stained for 15 min. After removing staining solution and washing in PBS twice, cells were imaged with microscope.

### Ethics statement

Primary peripheral blood samples including PDAC and normal subjects were collected from clinical residual blood samples with the waiver of informed consent from ethics approval (2022-0678). All experimental process was conducted under the permission of the Ethics and Scientific Committee of The Second Affiliated Hospital of Zhejiang University School of Medicine. All animal studies were conducted according to the national guidelines and were reviewed and confirmed by the ethics committee of Second Affiliated Hospital, School of Medicine, Zhejiang University (2023-152).

### Statistical analysis

All data is plotted as mean ± standard error of the mean (SEM). All figures and statistical analyses (Pearson correlation, two-tailed t-test, log-rank test, Tukey’s multiple comparisons test, Dunn’s multiple comparisons test) were performed using Prism software (GraphPad 8, San Diego, CA). Sample size (n) for each statistical analysis was indicated in figure legend. Differences with p-value less than 0.05 were considered statistically significant. Differences are labeled as follows: ns p>0.05, * p < 0.05; ** p < 0.01; *** p< 0.001; **** p < 0.0001.

## Funding

This work was supported by Weben fund (491020-15170C).

## Author contributions

R.Z.X conceived the study, initiated, designed, and supervised the experiments. R.Z.X, S.W.Z and X.X.G wrote the manuscript. S.W.Z, M.Y.L, Z.X.W, and X.Z.Z performed experiments. X.X.G conceived the study, and supervised the experiments.

## Diversity, equity, ethics, and inclusion [optional]

We welcome statements pertaining to diversity, equity, ethics, and inclusion, as outlined in our Editorial Policies.

## Competing interests

The authors declare no conflict of interest.

## Data and materials availability

All data are available in the main text or the supplementary materials.

## Notes

### Competing Interest Statement

The authors have declared no competing interest.

