## Supplementary figures 1-11 for "In vivo vaccine generation with a topical small molecule cocktail to eradicates AML and PDAC"

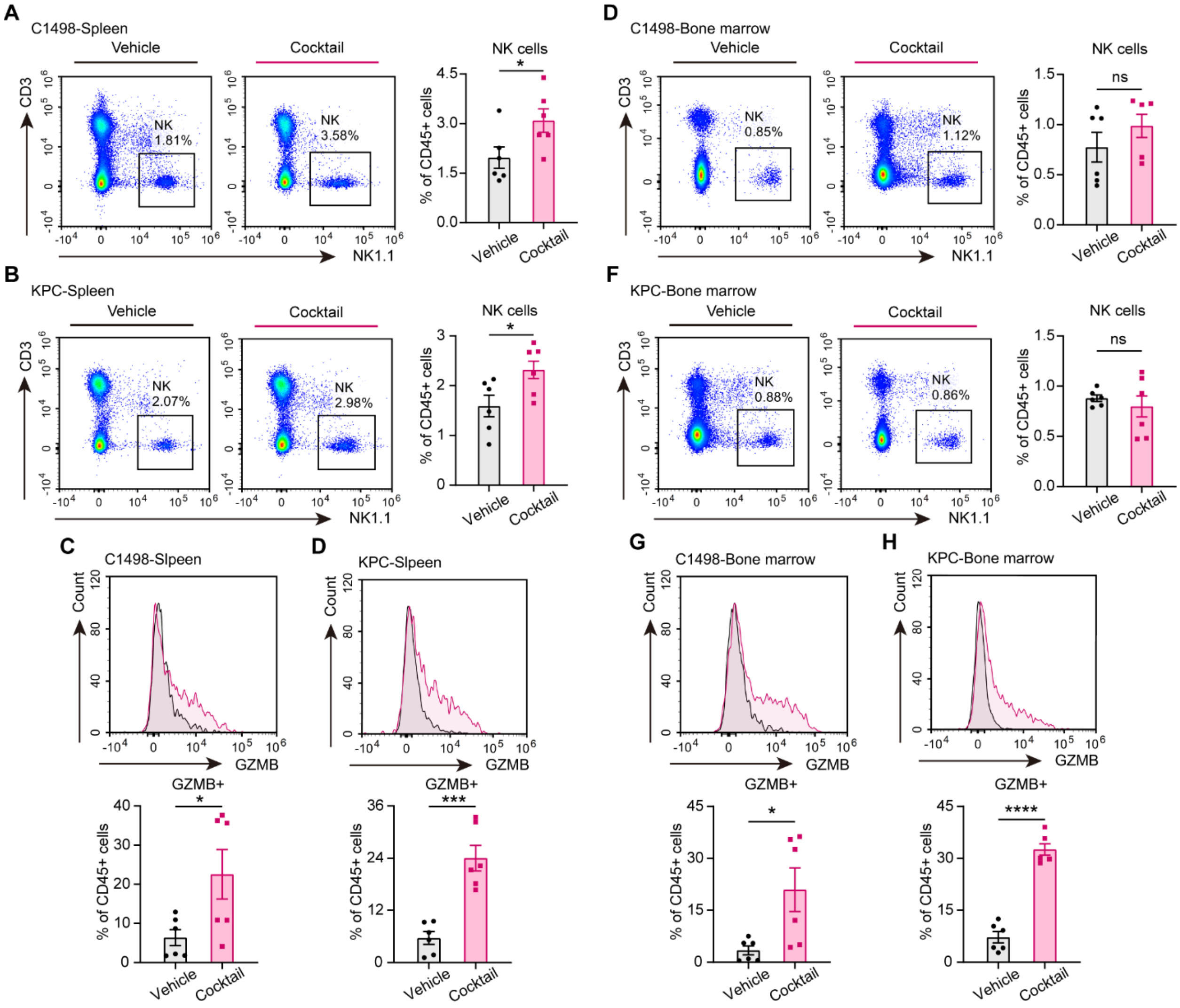


**Fig. S1.**

**Topical cocktail expands functional NK cells in spleen and bone marrow in mouse models.** **A. B.** Representative flow cytometry graphs and quantification of percentages of NK cells in $CD45^+$ cells in spleen of C1498 (**A**) and KPC (**B**) xenograft C57 mice on day7 after vehicle or vaccine treatment (n=6). **C. D.** Representative flow cytometry graphs (left) and quantification of percentages of $GZMB^+$ cells (right) in NK cells in spleen of C1498 (**C**) and KPC (**D**) xenograft C57 mice on day7 after vehicle or vaccine treatment (n=6). **E. F.** Representative flow cytometry graphs and quantification of percentages of NK cells in $CD45^+$ cells in bone marrow of C1498 (**E**) and KPC (**F**) xenograft C57 mice on day7 after vehicle or vaccine treatment (n=6). **G. H.** Representative flow cytometry graphs and quantification of percentages of $GZMB^+$ cells in NK cells in spleen of C1498 (**G**) and KPC (**H**) xenograft C57 mice on day7 after vehicle or vaccine treatment (n=6). Gating strategy is depicted in Supplemental Figure 2**F**. All error bars represent means ± s.e.m. Statistical significance was determined by the two-tailed t-test. * p<0.05, *** p<0.001, **** p<0.0001, ns means no significance.

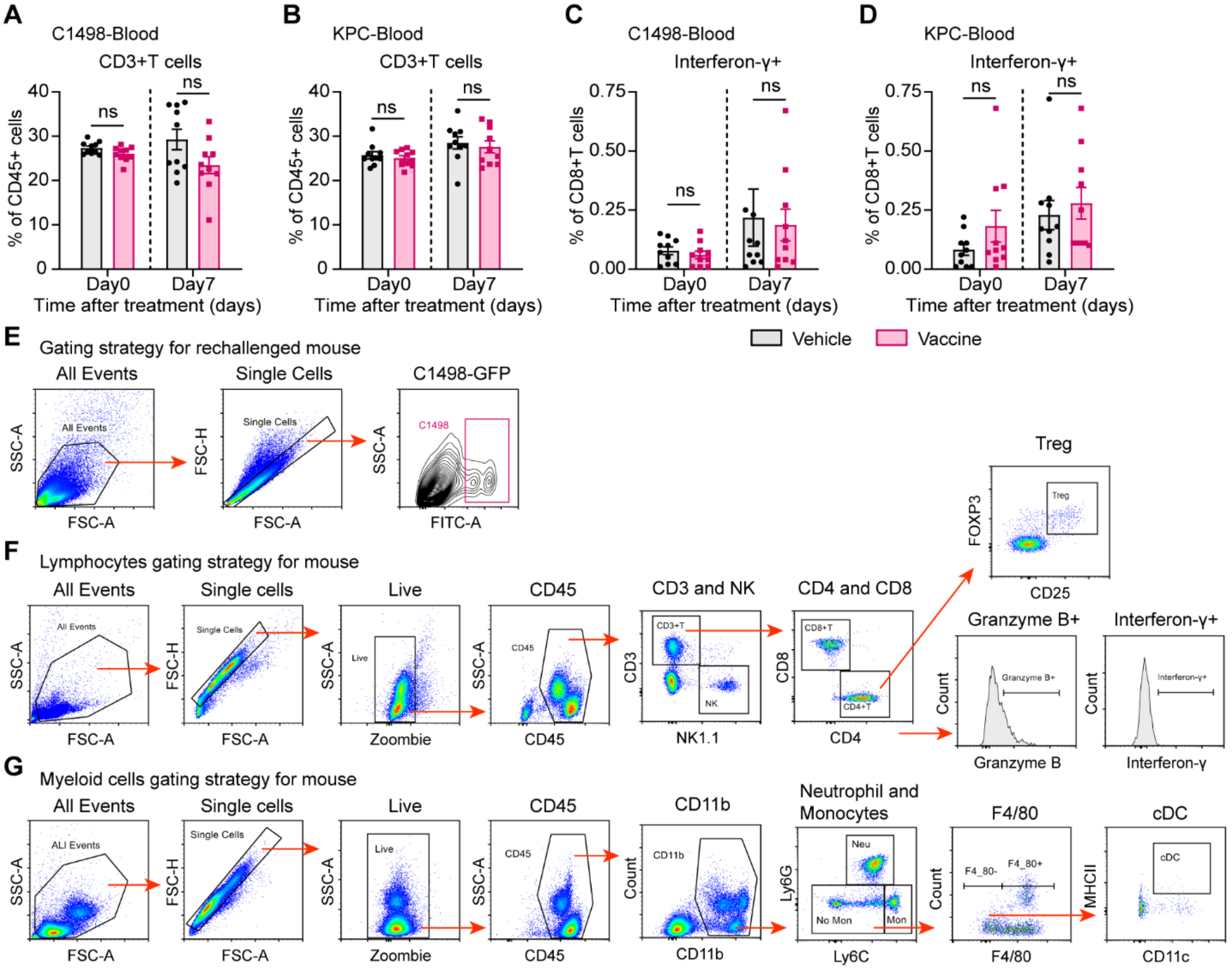


**Fig. S2.**

**Topical cocktail does not affect CD3$^+$T cells and IFNγ$^+$CD8$^+$ cells in circulating blood in mouse models. A. B.** Quantitative analysis of the percentage of CD3$^+$T cells in CD45$^+$ cells in blood of C1498 (**A**) and KPC (**B**) xenograft C57 mice on day0 and 7 after vehicle or vaccine treatment (n=10). **C. D.** Quantitative analysis of the percentage of Interferon-γ$^+$ cells in CD8$^+$T cells in blood of C1498 (**C**) and KPC (**D**) xenograft C57 mice on day0 and 7 after vehicle or vaccine treatment (n=10). **E.** Gating strategy for flow cytometry analysis of C1498-GFP cells from bone marrow, spleen and liver of rechallenged mice. **F.** Gating strategy for flow cytometry analysis of lymphocytes from blood, spleen and bone marrow of C1498 and KPC xenograft C57 mice. **G.** Gating strategy for flow cytometry analysis of myeloid cells from blood, spleen and bone marrow of C1498 and KPC xenograft C57 mice. All error bars represent means ± s.e.m. Statistical significance was determined by the two-tailed t-test, ns means no significance.

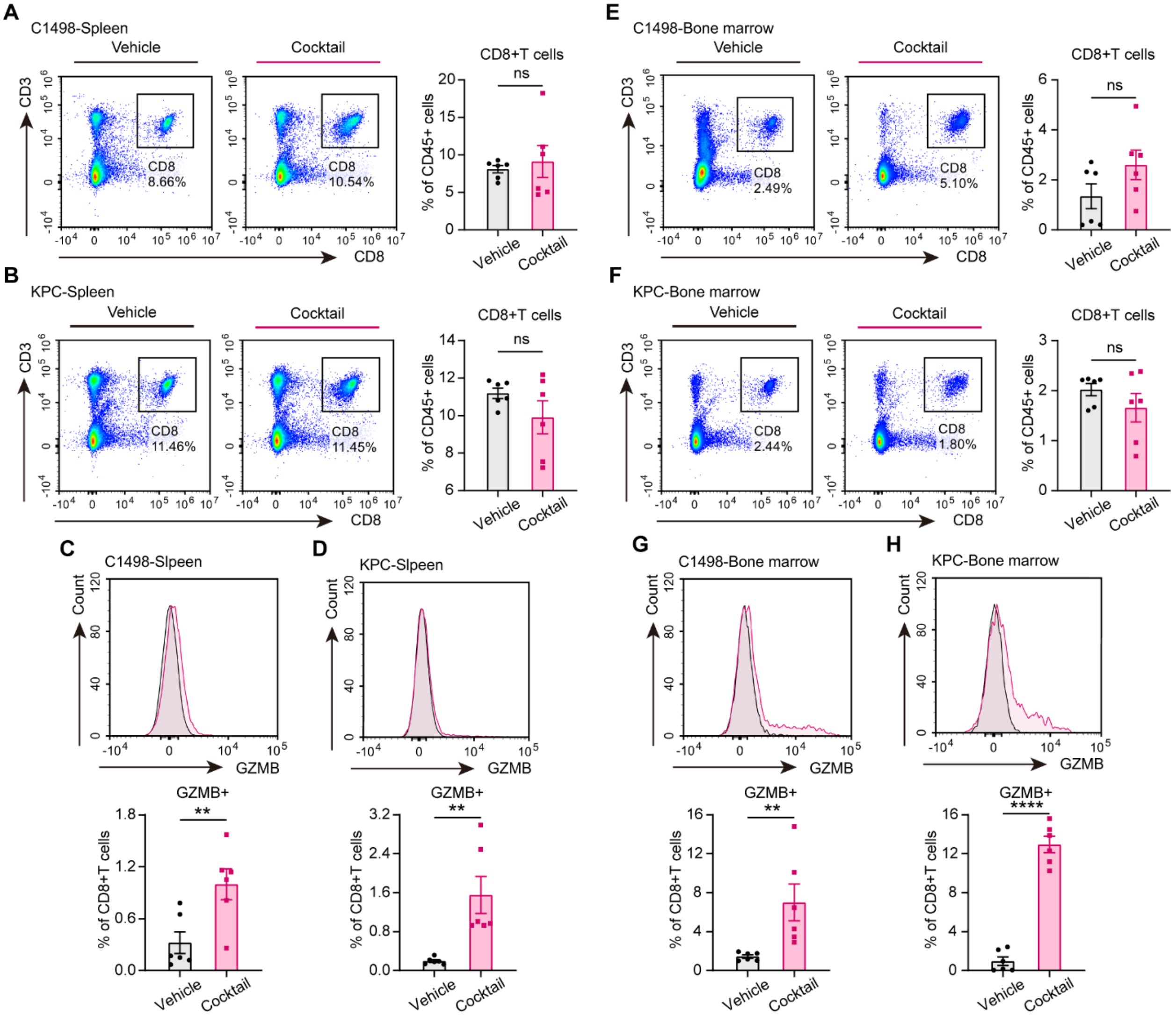


**Fig. S3.**

**Topical cocktail does not affect CD8$^+$T cells in spleens and bone marrow in mouse models.** **A. B.** Representative flow cytometry graphs and quantification of percentages of CD8$^+$T cells in CD45$^+$ cells in spleen of C1498 (**A**) and KPC (**B**) xenograft C57 mice on day7 after vehicle or vaccine treatment (n=6). **C. D.** Representative flow cytometry graphs and quantification of percentages of GZMB$^+$ cells in CD8$^+$T cells in spleen of C1498 (**C**) and KPC (**D**) xenograft C57 mice on day7 after vehicle or vaccine treatment (n=6). **E. F.** Representative flow cytometry graphs and quantification of percentages of CD8$^+$T cells in CD45$^+$ cells in bone marrow of C1498 (**E**) and KPC (**F**) xenograft C57 mice on day7 after vehicle or vaccine treatment (n=6). **G. H.** Representative flow cytometry graphs and quantification of percentages of GZMB$^+$ cells in CD8$^+$T cells in spleen of C1498 (**G**) and KPC (**H**) xenograft C57 mice on day7 after vehicle or vaccine treatment (n=6). Gating strategy is depicted in Supplemental Figure 2**F**. All error bars represent means ± s.e.m. Statistical significance was determined by the two-tailed t-test. ** $p<0.01$, **** $p<0.0001$, ns means no significance.

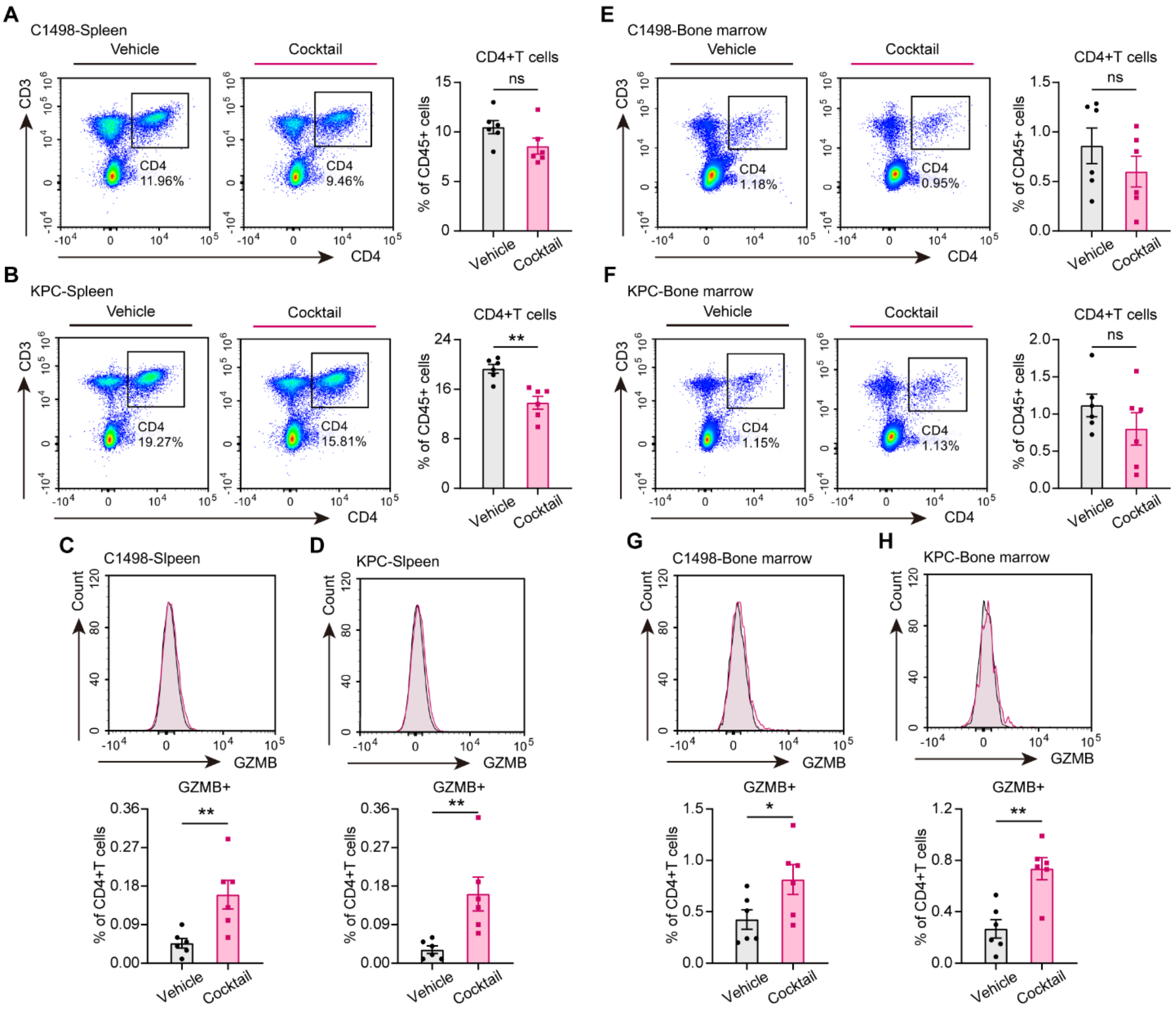


**Fig. S4.**

**Effect of topical cocktail on CD4 and CD4$^+$CTL cells in spleen and bone marrow in AML and PDAC mouse models. A. B.** Representative flow cytometry graphs and quantification of percentages of CD4$^+$T cells in CD45$^+$ cells in spleen of C1498 (**A**) and KPC (**B**) xenograft C57 mice on day7 after vehicle or vaccine treatment (n=6). **C. D.** Representative flow cytometry graphs and quantification of percentages of GZMB$^+$ cells in CD4$^+$T cells in spleen of C1498 (**C**) and KPC (**D**) xenograft C57 mice on day7 after vehicle or vaccine treatment (n=6). **E. F.** Representative flow cytometry graphs and quantification of percentages of CD4$^+$T cells in CD45$^+$ cells in bone marrow of C1498 (**E**) and KPC (**F**) xenograft C57 mice on day7 after vehicle or vaccine treatment (n=6). **G. H.** Representative flow cytometry graphs and quantification of percentages of GZMB$^+$ cells in CD4$^+$T cells in spleen of C1498 (**G**) and KPC (**H**) xenograft C57 mice on day7 after vehicle or vaccine treatment (n=6). Gating strategy is depicted in Supplemental Figure 2**F**. All error bars represent means ± s.e.m. Statistical significance was determined by the two-tailed t-test. ** p<0.01, **** p<0.0001, ns means no significance.

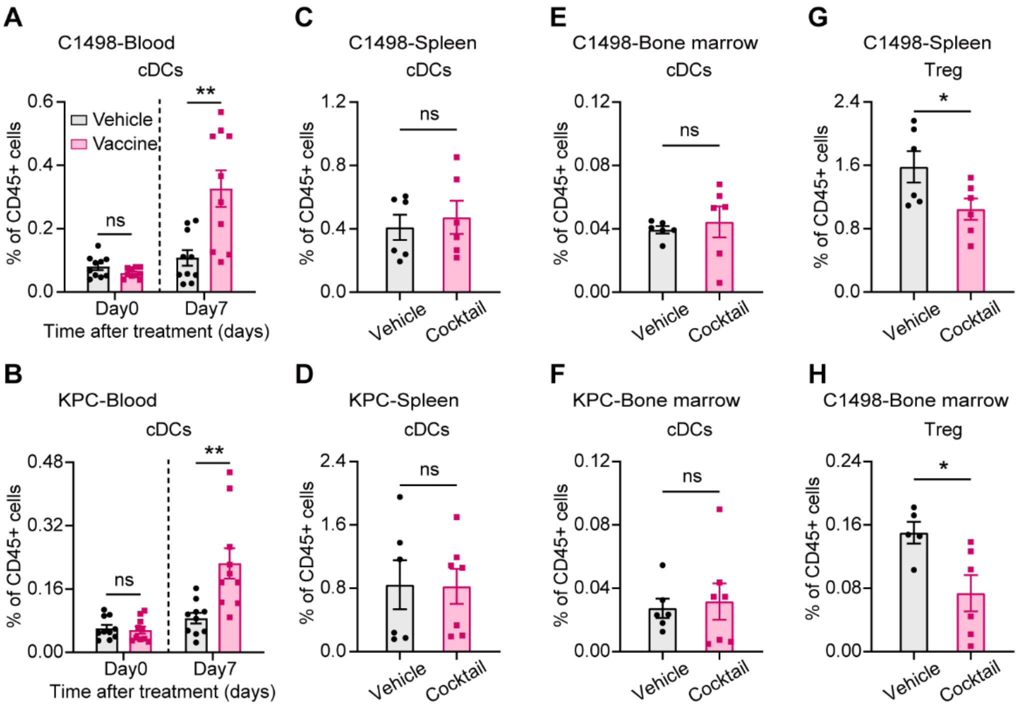


**Fig. S5.**

**The in vivo vaccine expands DC cells and reduces Treg T cells in AML and PDAC mouse models. A. B.** Quantification of percentages of cDCs in $CD45^+$ cells in blood of C1498 (**A**) and KPC (**B**) xenograft C57 mice on day0 and 7 after vehicle or vaccine treatment (n=10). **C-F.** Quantification of percentages of cDCs in $CD45^+$ cells in spleen (**C, D**) and bone marrow (**E, F**) of C1498 (**c, e**) and KPC (**D, F**) xenograft C57 mice on day0 and 7 after vehicle or vaccine treatment (n=6). **G. H.** Quantification of percentages of treg cells in $CD45^+$T cells in spleen (**G**) and bone marrow (**H**) of C1498 xenograft C57 mice on day7 after vehicle or vaccine treatment (n=6). Gating strategy is depicted in Supplemental Figure 2**F**, **G**. All error bars represent means ± s.e.m. Statistical significance was determined by the two-tailed t-test. * $p< 0.05$, ** $p<0.01$, ns means no significance.

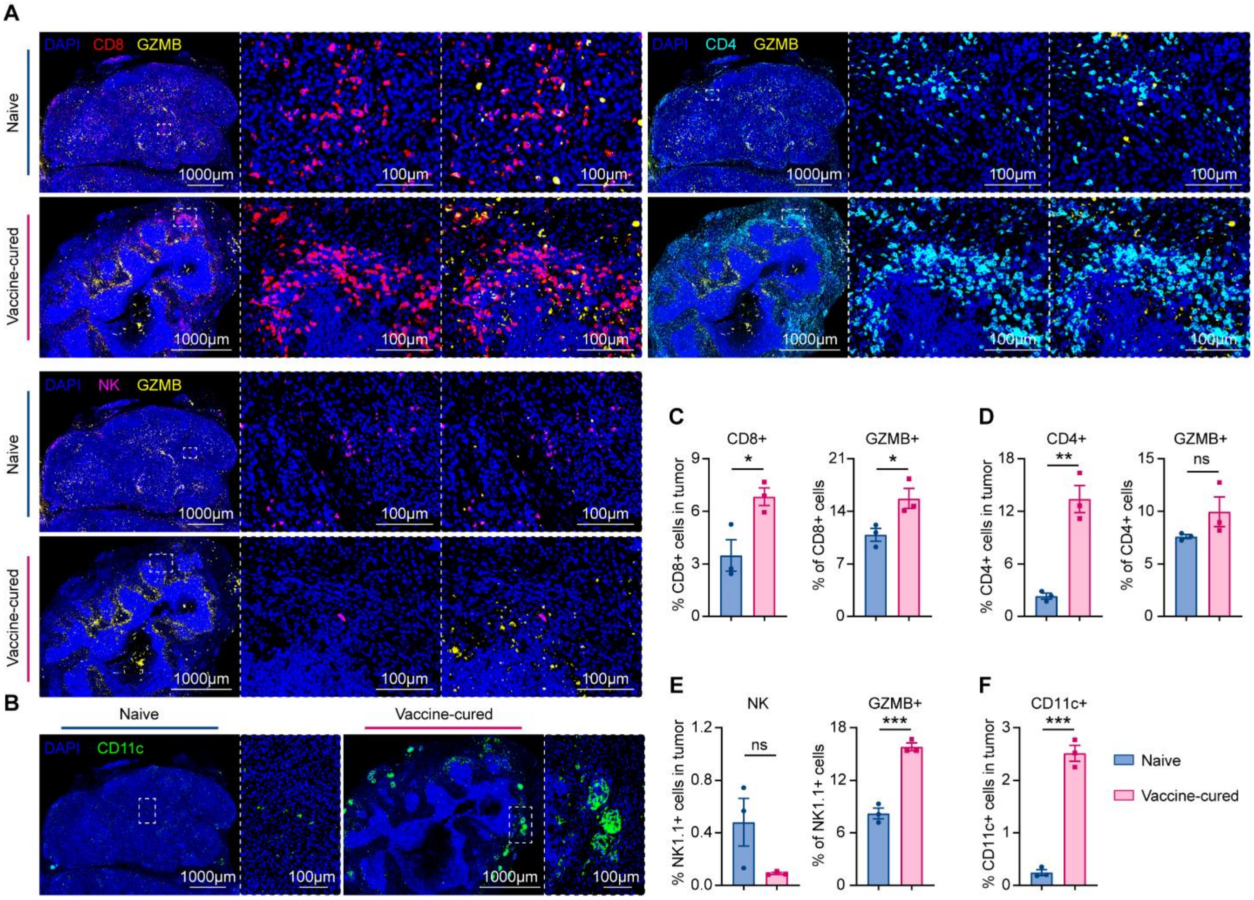


**Fig. S6.**

**The in vivo vaccine generates a super immune cell network in TEM in rechallenged mice.** **A. B.** Representative and the corresponding local enlarged images of GZMB+ (yellow) $CD8^+$ (Red) T cells, $GZMB^+$ (yellow) $CD4^+$ (Cyan) T cells, $GZMB^+$ (yellow) NK (Magenta) cells and CD11c (Green) staining of tumors from rechallenged mice, counterstained with DAPI (Blue). **C-F.** Quantification of $CD8^+$ T, $GZMB^+$ $CD8^+$ T (**C**), $CD4^+$ T, $GZMB^+$ $CD4^+$ T (**D**), NK, $GZMB^+$ NK (**E**) and $CD11c^+$ (**F**) cells in tumor (n=3). All error bars represent means ± s.e.m. Statistical significance was determined by the two-tailed t-test. * $p<0.05$, ** $p<0.01$, *** $p<0.001$, ns means no significance.

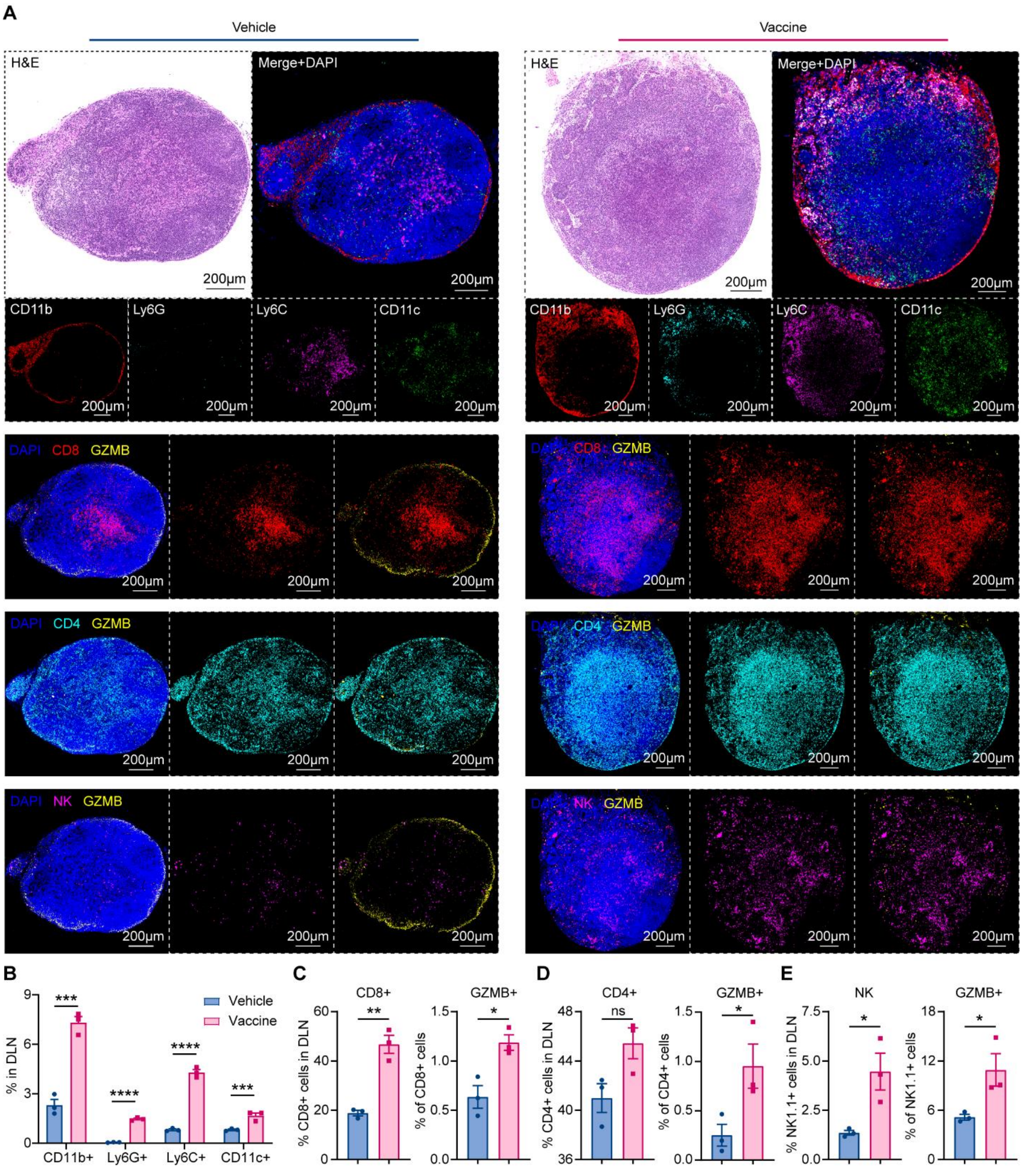


**Fig. S7.**

**The in vivo vaccine generates a super immune cell network in draining lymph nodes in PDAC mouse models. A.** Representative images of H&E, myeloid, $GZMB^+$ (yellow) $CD8^+$ (Red) T cells, $GZMB^+$ (yellow) $CD4^+$ (Cyan) T cells, and $GZMB^+$ (yellow) NK (Magenta) cells staining of draining lymph nodes (DLNs) from mice treated of vehicle or vaccine, counterstained with DAPI (Blue). Myeloid cells included $CD11b^+$ (Red), $Ly6G^+$ (Cyan), $Ly6C^+$ (Magenta), $CD11c^+$ (Green) cells. **B-E.** Quantification of myeloid (**B**), $CD8^+$ T, $GZMB^+$ $CD8^+$ T(**C**), $CD4^+$

T, GZMB$^+$ CD4$^+$ T (**D**), NK and GZMB$^+$ NK (**E**) cells in DLN (n=3). All error bars represent means ± s.e.m. Statistical significance was determined by the two-tailed t-test. * $p<0.05$, ** $p<0.01$, *** $p<0.001$, **** $p<0.0001$, ns means no significance.

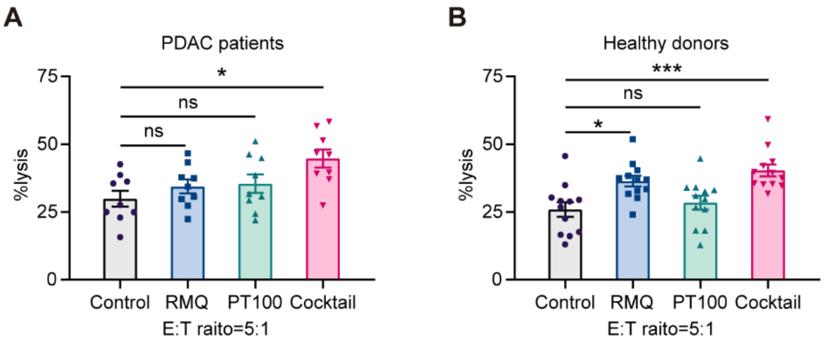


**Fig. S8.**

**Cocktail enhances PBMCs anti-tumor activity from PDAC patients and healthy donors. A. B**. Lysis of MIA-paca2-luciferase tumor cell lines (Target cells, T) by PBMCs (Effector cells, E) from PDAC patients (**A**) or healthy donors (**B**) was determined by luciferase-based cytotoxicity assay after 48h co-culture with RMQ (0.5μM), PT100 (0.5μM) or cocktail at the E:T ratio of 5: 1. All error bars represent means ± s.e.m. Statistical significance was determined by one-way ANOVA. * $p<0.05$, *** $p<0.001$, ns means no significance.

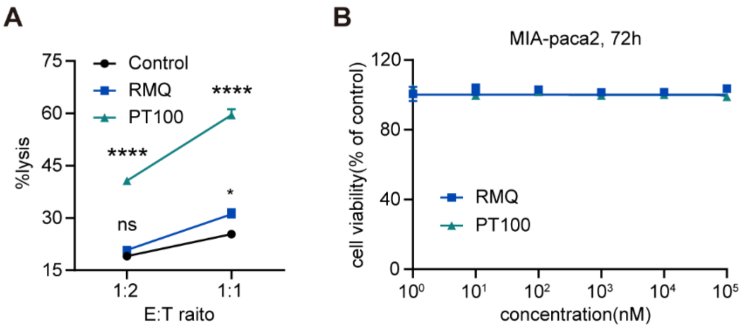


**Fig. S9.**

**Cocktail enhances NK cell anti-tumor activity. A.** Lysis of MIA-paca2-luciferase tumor cell lines (Target cells, T) by NK92 cells (Effector cells, E) was determined by luciferase-based cytotoxicity assay after 24h co-culture with RMQ (0.5μM) or PT100 (0.5μM) at the indicated E:T ratios. **B.** The rate of cell viability of the PDAC MIA-paca2 cells treated with RMQ or PT100 was measured by MTT assay after 72h. All error bars represent means ± s.e.m. Statistical significance was determined by one-way ANOVA. * $p<0.05$, **** $p<0.0001$, ns means no significance.

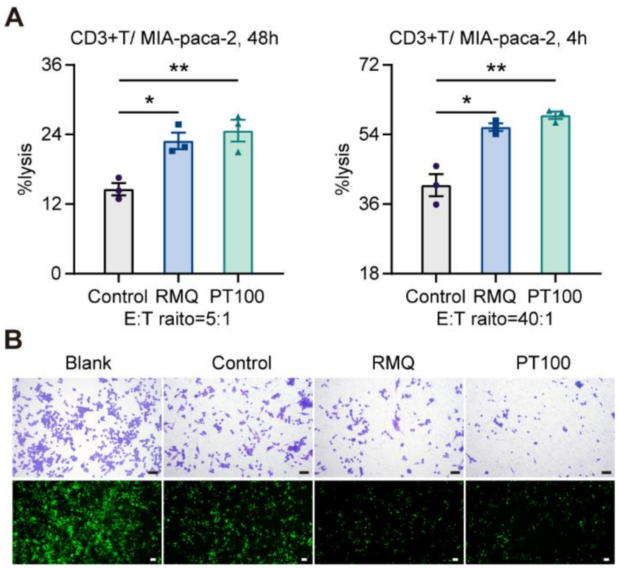


**Fig. S10.**

**small molecule cocktail enhances CD3$^+$T cell anti-tumor activity from patients with PDAC.** **A.** Lysis of MIA-paca2-luciferase tumor cell lines (Target cells, T) by CD3$^+$ T cells (Effector cells, E) was determined by luciferase-based cytotoxicity assay after 4h (left) or 48h (right) co-culture with RMQ (0.5μM) or PT100 (0.5μM) at the E:T ratio of 5:1. **B.** Representative images of tumor lysis ability of CD3$^+$ T cells. CD3$^+$ T cells were incubation with MIA-paca2-GFP cells after treated with RMQ (0.5μM) or PT100 (0.5μM) after 48h at the E:T ratio of 5:1. The tumor lysis ability was detected by crystal violet assay (upper) and GFP scanning (below). Scale bars: 40 μm. All error bars represent means ± s.e.m. Statistical significance was determined by one-way ANOVA. * $p<0.05$, ** $p<0.01$.

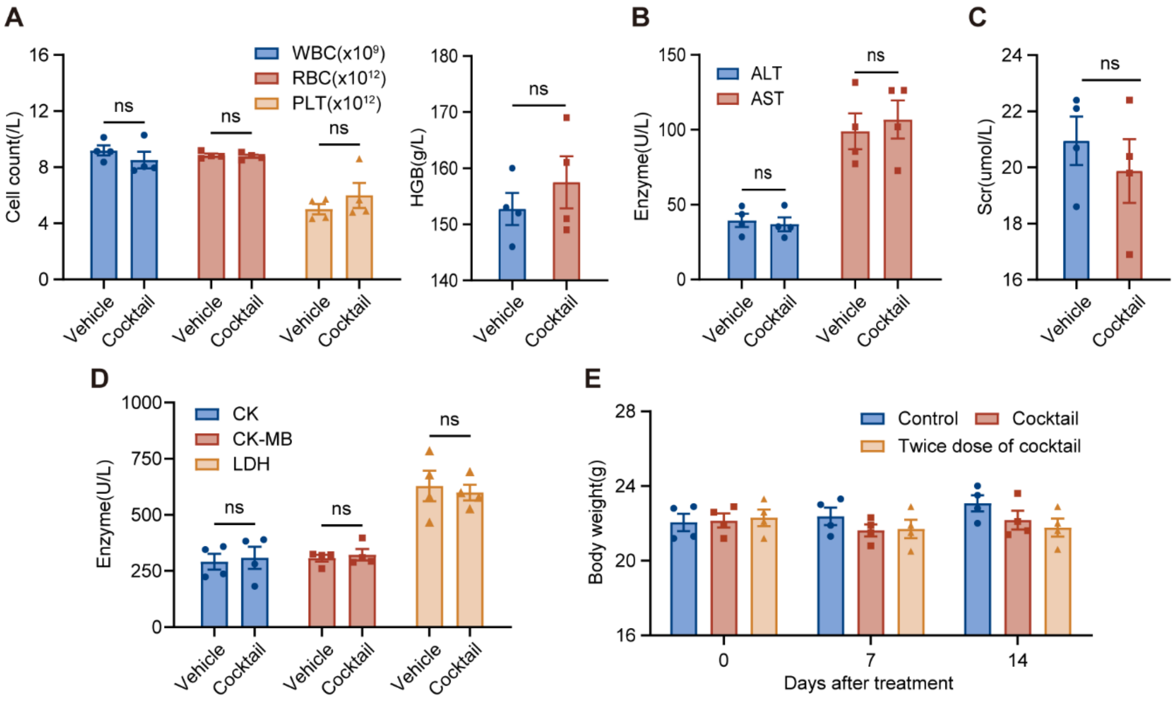


**Fig. S11.**

**Safety assessment of in vivo vaccine in vivo. A-D.** Safety-associated indicators of in vivo vaccine, including blood cell counts (**A**), HGB (**A**), liver enzyme (**B**), serum creatinine (**C**), and myocardial enzymes (**D**) were evaluated after treatment with vehicle or cocktail for once a week for 3 weeks (n=4). **E.** Body weight of C57 mice after treatment with vehicle, cocktail, or twice dose of cocktail on day0, 7, and 14 (n=4). WBC: white blood cell; RBC: red blood cell; PLT: platelet count; HGB: hemoglobin; ALT: alanine aminotransferase; AST: aspartate transaminase; Scr: serum creatinine; CK: creatine kinase; CK-MB: creatine kinase isoenzymes; LDH: lactate dehydrogenase. All error bars represent means ± s.e.m. Statistical significance was determined by the two-tailed t-test, ns means no significance.
